# Corroborating age with oxygen isotope profiles in otoliths: consequences for estimation of growth, productivity and management reference points in northern pike (*Esox lucius*)in the southern Baltic Sea

**DOI:** 10.1101/2023.02.01.526588

**Authors:** Timo D. Rittweg, Clive Trueman, Elias Ehrlich, Michael Wiedenbeck, Robert Arlinghaus

**Affiliations:** Leibniz Institute of Freshwater Ecology and Inland Fisheries (IGB), Müggelseedamm 310, 12587 Berlin, Berlin, Germany; School of Ocean and Earth Science, University of Southampton Waterfront Campus, European Way, SO143ZH Southampton, UK; German Research Center for Geosciences (GFZ) Potsdam, Telegrafenberg, 14473 Potsdam, Brandenburg, Germany; Division of Integrative Fisheries Management, Faculty of Life Sciences, Humboldt-Universität zu Berlin, Invalidenstr. 42, 10115 Berlin, Berlin, Germany

**Keywords:** Baltic Sea, coastal waters, brackish, aging, ageing, von Bertalanffy, Bayesian growth modeling, yield, d18O, stable isotopes, scales, secondary ion mass spectrometry

## Abstract

Accurate and precise age estimates are crucial for assessing the life-history of fish and providing management advice for fisheries, but age validation studies remain rare or absent in many species. Aging from scales is common, as it is non-lethal, but potential for underaging old fish exists. Using 85 northern pike (*Esox lucius*) collected from the southern Baltic Sea in Germany as a model, we corroborated age readings based on annual cycles of oxygen isotopes (*δ*^18^*O*) in otoliths to infer the timing and validity of growth, so as to compare results with visual age estimations from scales and otoliths. Otoliths were accurate and precise, while age readings from scales systematically underestimated the age of old pike. Fitting population-level von Bertalanffy growth models to the size-at-age data estimated via *δ*^18^*O*-profiling, otoliths or scales revealed a larger terminal length (*L*_∞_) and a lower body growth coefficient *k* in scale-aged fish compared to otolith and corroborated age data. Populating an age-structured model with structure-specific growth model parameters demonstrated that the maximum sustainable yield (*MSY*) was estimated to be about 37% lower using scale-informed growth models relative to growth models fitted to corroborated and otolith-based size-at-age data. Thus, pike populations assessed and modeled based on scale age readings might appear less productive than they really are. Using scale-based ages to inform management regulations may therefore result in too conservative management and lost biomass yield, while instilling unrealistic angler expectations as to the trophy potential of the fishery.

## 1. Introduction

Estimation of exploitation status and resilience of fish stocks, together with ecological assessment of factors affecting life history, population and community dynamics, form the core of evidence-based fisheries management. Most life-history traits of fish have a time dimension, e.g., age-at-maturation, growth rate, mortality or longevity (Ahrens et al., 2020; Beverton and Holt, 1957; Denechaud et al., 2020). Counting periodic growth increments that form on calcified hard structures remains the most common approach to aging fish (Campana et al., 1995; Denechaud et al., 2020). Aging is central to most fisheries assessment methods (Maceina et al., 2007; Quist et al., 2012; Spurgeon et al., 2015), especially when catch-at-age time series are used (Fournier and Archibald, 1982), but also when relying on data-poor assessment alternatives (Lorenzen et al., 2016) or simulation models which need age structure, natural mortality rate or growth rate estimates as inputs (Beverton and Holt, 1957; Pierce et al., 2003). Errors in age estimation can thus lead to ill-informed reference points and mismanagement (e.g., incorrect harvest regulations), which may have serious consequences for fish populations (Reeves, 2003; Tyszko and Pritt, 2017; Yule et al., 2008). Information on accuracy and precision of aging structures and methods along with the range of ages on which they are applicable are a prerequisite for robust conclusions on life history and population dynamics of a managed fish species. These data are particularly rare in inland and small-scale coastal fisheries that lack resources for annual monitoring (Beard et al., 2011; Brownscombe et al., 2019; Midway et al., 2016; Post et al., 2002). One example where severe data gaps exist are coastal fisheries in the southern Baltic Sea, such as the fisheries targeting the freshwater species northern pike (*Esox lucius*, Linnaeus 1758) in brackish environments (Gemert et al., 2022).

Baltic Sea pike stocks have traditionally been exploited by commercial fisheries, but in recent years a strong recreational fishery has developed that often has different objectives to commercial fisheries, e.g., favoring the catch of trophy fish or catch rate over biomass yield (Ahrens et al., 2020; Arlinghaus et al., 2019). Over the last decade, pike stocks have declined in many areas of the Baltic (Berggren et al., 2021; Bergström et al., 2022; Lehtonen et al., 2009; Olsson, 2019; Tibblin et al., 2015). To align fishing mortality rates with shifts in resource abundance, assessment of contemporary growth rates and derivation of fisheries reference points that suit commercial and recreational fisheries is important (Ahrens et al., 2020). This, in turn, demands an assessment of possible sources of bias in aging procedures for northern pike in the Baltic and testing of the effects aging errors have on growth modelling and subsequent derivation of management reference points.

Aging errors are classified into process errors and interpretation errors (Beamish and McFarlane, 1995; Campana, 2001; Maceina et al., 2007). Process errors result from aging structures which do not form true annual marks, or annual marks are formed but are not visually discernible with a given aging method, preventing a reliable estimation of age. Process errors are thus errors of accuracy, and are usually biased towards under- or overestimation of age, while interpretation errors are a result of poor reproducibility of age estimates due to individual subjectivity, and can be either biased or random (Beamish and McFarlane, 1995; Campana, 2001; Maceina et al., 2007). Both error types can have multiple underlying explanations, such as changing environmental conditions, ontogenetic transitions (e.g., change in prey type) and slow growth rates that form subannual checks on hard structures that are misinterpreted as annual marks (annuli) (Weyl and Booth, 1999), or blur true annuli (Casselman, 1990; Weyl and Booth, 1999; Whiteman et al., 2004). Underestimation of age can lead to stocks appearing faster growing and more productive than they really are (Beamish, 1987; Lai and Gunderson, 1987), leading to overly optimistic estimates of growth and positive bias in mortality rate (Campana, 1990; Chilton and Beamish, 1982; Lai and Gunderson, 1987; Van Den Broek, 1983). Overestimation of age is less common and perhaps less consequential in fisheries models (Lai and Gunderson, 1987), but has been found to affect estimates of stock resilience to harvest in some cases (e.g., Bertignac and Pontual (2007)). To mediate such errors in advance of deriving management suggestions from such models, age validation or age corroboration efforts are a necessary precaution (Beamish and McFarlane, 1995; Maceina et al., 2007; Spurgeon et al., 2015).

Age validation efforts of fish aim at confirming the temporal scale at which visual growth marks are deposited in a calcified structure (Beamish and McFarlane, 1983). They can be divided into three broad categories (Campana, 2001). The first category involves techniques validating the absolute (i.e., true) age of individuals, such as known-age mark recapture (Bruch et al., 2009; Crane et al., 2020; Hamel et al., 2014) or bomb-radiocarbon dating (Bruch et al., 2009; Casselman et al., 2019). The second approach validates the periodicity of growth mark deposition, such as the release and recapture of chemically (e.g., tetraoxycycline) marked wild-caught fishes (Brown and Gruber, 1988; Laine et al., 1991). The third approach does not validate ages per se, but instead aims at corroborating age estimates of a sample of unmarked fish of unknown age (Kimura et al., 2006; Maceina et al., 2007). Age corroboration is advisable when no validated ages or known-age fish are available, and the species in question is not particularly long-lived (Heimbrand et al., 2020; Hüssy et al., 2020a; Kimura et al., 2006; Van Oosten, 1923). This is the case for Baltic pike, for which, to our knowledge, no age validation or known age fish exist so far.

One approach for corroborating a set of age estimates is studying the periodicity of seasonal variation through elemental cycles recorded in fish otoliths (Heimbrand et al., 2020; Hüssy et al., 2020a; Kastelle et al., 2017; Weidman and Millner, 2000). Otoliths record information from ambient water by incorporating elements through continuous precipitation of aragonite mineral (Darnaude et al., 2014; Darnaude and Hunter, 2018; Hüssy et al., 2020b; Kerr et al., 2007; Matta et al., 2013). A promising approach of inferring periodicity through elemental cycles is the ratio of oxygen isotopes. The relative abundance of two stable isotopes of oxygen, ^18^O to ^16^O, notated as *δ*^18^*O* values,varies as a function of the isotopic composition and temperature of the ambient water and has been demonstrated to be a reliable thermal marker for aquatic species (Darnaude et al., 2014; Matta et al., 2013; Morissette et al., 2020; R. Naylor et al., 2007), including fish (Hoie and Folkvord, 2006; Kastelle et al., 2017; Weidman and Millner, 2000). As pike prefer shallow habitats, and the species has its main distribution range in temperate regions (Casselman, 1996), pike commonly are exposed to a strong seasonal thermal range, expected to translate into well-developed seasonal *δ*^18^*O*-cycles in otoliths. This notion is supported by Gerdeaux and Dufour (2012), who demonstrated that *δ*^18^*O* values follow a strong seasonal cycle for pike in Lake Annecy, France. Methodological advancements over recent years allow the determination of *δ*^18^*O* values at high precision (< 0.1‰ standard deviation) and high spatial resolution (~40 *μm*) using secondary ion mass spectrometry (SIMS) transects on otoliths (Matta et al., 2013; Morissette et al., 2020). This approach also bears the advantage of measuring isotopic abundances on a standardized transect from core to margin, without prior interpretation of annual structures.

Age estimates of pike have been performed with a multitude of calcified structures, e.g., otoliths (Oele et al., 2015), scales (Anwand, 1969; Frost and Kipling, 1959), cleithra (Casselman, 1996, 1974), metapterygoid bones (Sharma and Borgstrom, 2007), pelvic fin rays (Babaluk and Craig, 1990) and opercula (Berggren, 2019). While otoliths were described by previous work as a precise aging structure causing minimal among-reader bias in pike (Blackwell et al., 2016; Oele et al., 2015), considerable debate exists on whether scales provide reliable age readings. Laine et al. (1991) report scales to be accurate up to an age of 10 years using tetraoxycycline marked cleithra as a reference structure, Pagel et al. (2015) found scale age estimates of juvenile pike to agree well with age estimates from cleithra, and Anwand (1969) also reported precise aging results using scales. However, several other studies reported lack of accuracy and precision in age estimates on pike scales, such as Mann and Beaumont (1990) in a known age mark-recapture study, Frost and Kipling (1959) by comparing age estimates from pike scales and opercula and, more recently, Oele et al. (2015) and Blackwell et al. (2016), who both compared multiple calcified structures in terms of precision and reader bias. Because scales continue to be used in pike studies (Monk et al., 2021), further age evaluation approaches are warranted, especially in the Baltic Sea where aging from scales and modelling growth of pike using scale-derived size-at-age data has been common in the past (Dorow, 2004; Droll, 2022; Hegemann, 1958; Juncker, 1988).

Hard structures in pike from the Baltic Sea might also differ in accuracy and precision compared to freshwater pike, as the natural habitat of a fish species can influence the visual patterns on calcified hard structures (Heimbrand et al., 2020). For example, in Baltic cod (*Gadus morhua* Linnaeus, 1758) it has been hypothesized that complex hydrography along with migrations across salinity and temperature gradients lead to high variability in the visual properties of calcified structures, resulting in poor precision and low accuracy (Hüssy et al., 2016). Examples for habitat-specific variation in visual patterns on hard structures also exist for pike, e.g., in fish from Slepton Ley, United Kingdom (Sharma and Borgstrom, 2007), for which scales and operculae proved particularly difficult to read. Hence, the optimal aging structures may need to be adapted to the specific system under study, and their accuracy and precision should be assessed for the entire age range of interest (Beamish and McFarlane, 1983; Spurgeon et al., 2015).

The objective of our study was to examine the accuracy and precision of age estimates from otoliths and scales for 85 pike collected from brackish lagoons of the southern Baltic Sea in Germany. We derived corroborated age estimates from *δ*^18^*O* chronologies on otoliths, which were compared with visual age estimates from scales and otoliths. Using structure-specific size-at-age estimates, we calculated von Bertalanffy (Von Bertalanffy, 1938) growth curves, and compared the growth parameters between the aging structures. We then used an age-structured population model to examine how the growth parameter estimates translated into estimates of fisheries reference points, such as maximum sustainable yield (*MSY*),fishing mortality at maximum sustainable yield (*F_MSY_*), and biomass at maximum sustainable yield (*B_MSY_*). Furthermore, we calculated the optimal minimum-length limit arising from each growth model, considering multiple management objectives. We hypothesized that (1) structures used to estimate pike age differ strongly in their accuracy and precision compared to a corroborated reference age with scale-derived age estimates underestimating the age of old pike, (2) age estimates derived from varying aging structures significantly affect population-level von Bertalanffy growth rates, overestimating the terminal length in scale-read size-at-age data, and (3) the different growth parameters from growth curves affect model-based estimation of fisheries reference points and optimal harvest regulations.

## 2. Material and Methods

### 2.1. Sampling of pike

Eighty-five pike were caught between fall 2019 and spring 2021 in the three major lagoons (called Bodden) in the southern Baltic Sea around the Island of Rügen, Germany (Figure **1**): Northern Rügen Bodden Chain (NRB), western Rügen Bodden Chain (WRB) and the Greifswalder Bodden (GB). Additional fish were sampled from the freshwater tributaries Barthe and Peene Rivers, Sehrowbach, Ziese, Neuendorfer Hechtgraben (NHG) and Badendycksgraben (BKG). Fish were sampled with fyke nets, gill nets, rod and line fishing and electrofishing. Length of each fish at capture was determined to the nearest mm. A subsample of pike from the sample pool were chosen for secondary ion mass spectrometry (SIMS) analysis at the German Research Center for Geosciences (GFZ) in Potsdam, Germany. We attempted to sample fish from all major lagoons across the entire length-gradient and with a male to female ratio of 1:1. However, sampling limitations did not allow an equal sex ratio. We complemented the lagoon sample with pike collected from lagoon tributaries. Our aims were not to directly compare the sizes at ages among lagoons and tributaries, but to carry a representation of different environments and sexes into our final pool of samples. The final sample size was 24 individuals in WRB, 22 in NRB, 18 in GB and 24 individuals from freshwater streams. One fish was excluded from analysis due to a misplaced SIMS transect, two fish were excluded because no scales with a complete growth record were available, adding up to a final sample size of 85 fish.

**Figure 1:**
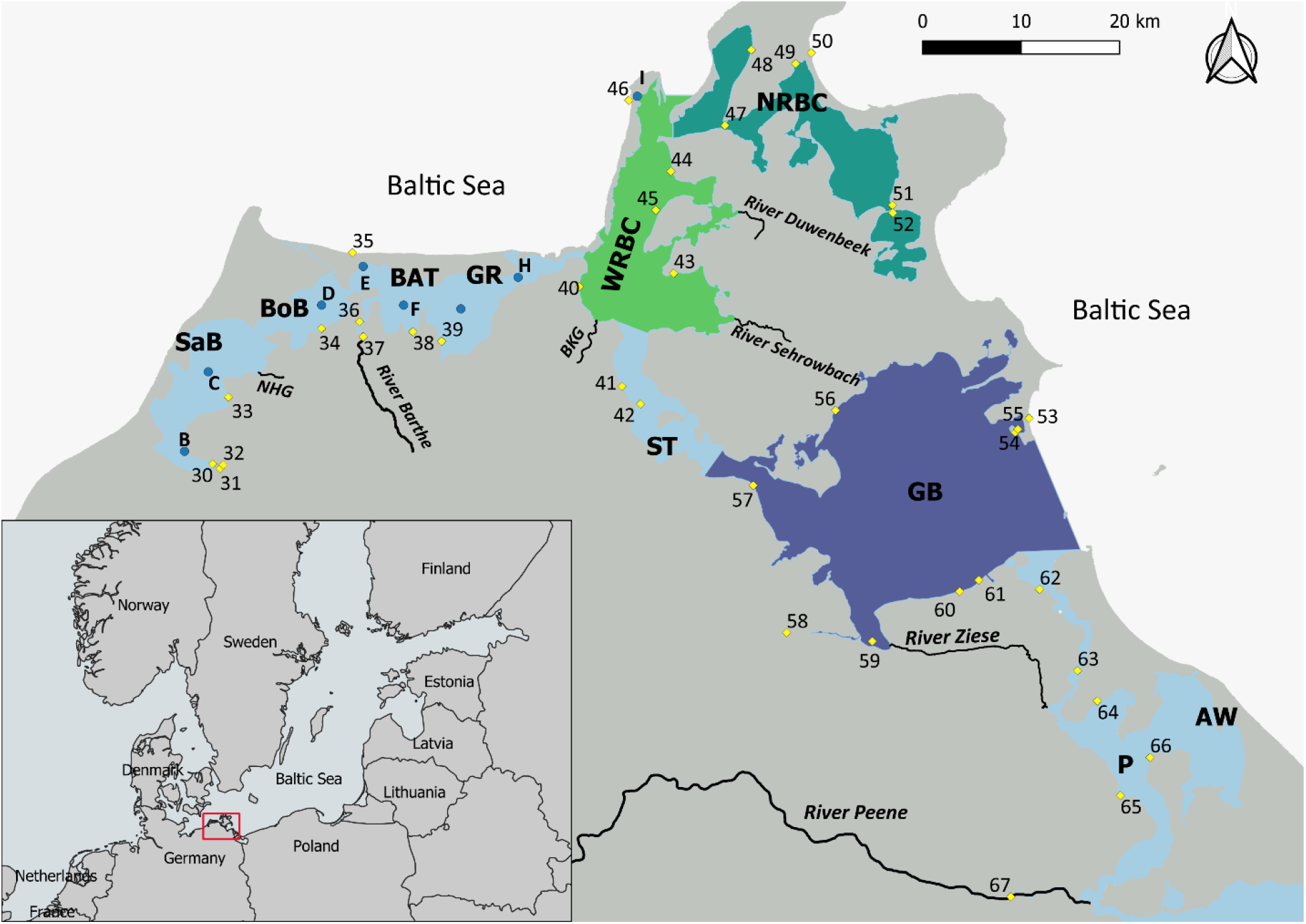
Study area around the island of Rügen in the southern Baltic sea, depicting the three major lagoon chains from which pike have been sampled for this study, along with the water isotope sampling points of Aichner et al. (2021). Yellow marks labelled with numbers indicate isotope transect points sampled in June 2019, March 2020 and July 2020, Blue marks labelled with letters indicate isotope timeseries stations, sampled from March 2020 until March 2021 weekly (for points D - H) or biweekly (for point I). NRBC: Northern Rügen Bodden Chain; WRBC: Western Rügen Bodden Chain; GB: Greifswalder Bodden; SaB: Saaler Bodden; BoB: Bodstedter Bodden; BAT: Barther Bodden; GR: Grabow; ST: Strelasund; P: Peenestrom; AW: Achterwasser; NHG: Neuendorfer Hechtsgraben; BKG: Badendycksgraben.

### 2.2. Scale processing

Scales were sampled from above the lateral line just anterior to the dorsal fin and placed in paper envelopes to dry for a minimum of three days to improve readability, then soaked in soap water, cleaned and placed on object slides labeled with the individual ID. A minimum of three scales per fish was mounted per slide. The slides were placed under a Leica MZ8 stereo microscope using a plan apochromat M objective and photographed using a Leica MC190 HD microscope camera. Images were contrast-enhanced in the Fiji distribution of ImageJ (Schindelin et al., 2012) via histogram equalization. Images were produced from three different scales for each fish for aging, and read randomly without knowing the size, origin, or sex of the fish.

### 2.3. Otolith processing

Sagittal otoliths were extracted, adhering remains of the labyrinth and mucosal tissue were removed, and otoliths were then transferred to 15 ml polypropylene tubes filled with ultrapure water (MilliQ, conductivity < 60 *μS* * cm^-1^). The tubes were placed in an ultrasonic waterbath and sonified for 5 min. Afterwards, the water was decanted and the tube filled with fresh ultrapure water, and the cleaning procedure was repeated. Otoliths were sonified a total of 3 times, then placed into acid-washed Eppendorff tubes. To avoid contamination with metal, otoliths were handled exclusively with pre-rinsed polypropylene tweezers after the cleaning procedure and were left to dry in a desiccator for 48 h. Once dry, the otoliths were glued to an object slide with Crystalbond and a section of 100 *μm* in width was cut out of the center of the otolith with an Isomet low speed saw (BUEHLER Ltd 11-1180). After cutting, the Crystalbond glue was dissolved in LC-MS grade Acetone (ROTISOLV ≥ 99.9 %). Thin sections were polished with 3000 and 5000 grit sandpaper and placed in dry acid-washed Eppendorf tubes, labelled and sent to the secondary ion microprobe facility at German Research Center for Geosciences Potsdam (GFZ) for further processing.

### 2.4. Secondary ion mass spectrometry of otolith *δ*^18^*O*

The otolith sections were embedded into SIMS sample mounts with Epofix epoxy resin, along with the UWC3 and IAEA603 calcite reference materials (Kozdon et al., 2009; IAEA 2016). The mount was polished until a surface quality of < 10 *μm* was reached. Surface quality was assessed via white light profilometry. Overview mosaic images with 2.5 X magnification of the mounts were produced via the stitching software of a Nikon Eclipse motorized optical microscope. Transects were marked digitally on the otolith sections in a straight line from the core to the distal edge along the longest available axis crossing all visible annuli. Transect lines were chosen to overshoot the otolith core to ensure the transect covered a complete line from core to edge. Sample mounts were sputter-coated with a 35 nm thick, high-purity gold film, placed in non-magnetic sample holders in a specially designed high-vacuum storage chamber of the Cameca 1280-HR secondary-ion mass spectrometer. Analyses were performed as point profiles with a step size of ~35 μm along the transects, with both reference materials being analyzed after each 10^th^ otolith determination. All *δ*^18^*O* analytical results were corrected for the determined instrumental mass fractionation, and were reported in ‰ relative to Vienna Standard Mean Ocean Water (VSMOW). Otolith *δ*^18^*O* values were converted to ‰ relative to Vienna Pee Dee Belemnite (VPDB) using the conversion equation described in Kim et al. (2015) for further analysis.

### 2.5. Otolith imaging

After SIMS *δ*^18^*O* data acquisition, the sample mounts were cleaned with ethanol and repolished with 300 nm Alpha alumina powder diluted with distilled water to remove the gold coating. The mounts were placed under a Nikon Eclipse motorized optical microscope and mosaic images of individual otoliths were taken under darkfield setting at 10 X magnification and stitched together by the stitching software of the Nikon Eclipse microscope. The resulting high-resolution images made it possible to visually discern the spots from the SIMS transects. The images were used both for acquiring otolith age estimates using visual readings and to perform one-to-one mapping of *δ*^18^*O* transects on the images.

### 2.6. Expected *δ*^18^*O* values in otolith aragonite

To assess whether observed *δ*^18^*O* values are following a seasonal pattern, we calculated theoretical *δ*^18^*O* values in a pike otolith using water *δ*^18^*O* values from a sampling campaign in our study area carried out by Aichner et al. (2021), measuring salinity, temperature and *δ*^18^*O* at a measuring buoy located in one of our sample areas (Vitter Bodden, WRB). Water samples were taken biweekly from March 2020 to March 2021 from the lagoon at the ferry harbor of Kloster on the island of Hiddensee west of Rügen (time series point **I**, Figure **1**). Temperature and salinity were measured for surface and bottom water with a Multi 3630 IDS device (WTW, Weilheim, Germany), equipped with a WTW TetraCon 925 electrical conductivity measuring cell. Water samples were transferred to the water isotope lab at Leibniz Institute of Freshwater Ecology and Inland Fisheries (IGB) in Berlin. Samples were filtered through 0.2 *μm* acetate filters prior to analysis. The *δ*^18^*O* values were measured using a Picarro (Santa Clara, CA, USA) L2130-i cavity ring-down spectrometer (CRDS). Isotope values and standard deviations were based on six replicate measurements from each sample, discarding the first three measurements to avoid memory effects. Three laboratory reference materials were analysed after every 24 samples (L, *δ*^18^*O*= −17.86‰, DEL *δ*^18^*O*= −10.03‰ and H, *δ*^18^*O*= −7.68), with a fourth reference material, M (*δ*^18^O = −7.68‰) used to assess drift and quality after every sixth sample. In-house reference materials were linked to primary reference materials VSMOW2 (Vienna Standard Mean Ocean Water 2), GRESP (Greenland Summit Precipitation) and SLAP2 (Standard Light Antarctic Precipitation 2) from the International Atomic Energy Agency in Vienna (IAEA). We modeled theoretical *δ*^18^O-values for a pike otolith (hereafter *δ*^18^*O_Otolith_*) with measured temperature and water *δ*^18^*O*-values (hereafter *δ*^18^*O_Water_*). As temperatures measured on shore closely tracked measurements from the open lagoon (one-way ANOVA, F(1,46) = 0.12, *p* = 0.73) and surface temperatures closely followed bottom temperatures (one-way ANOVA, F(1,272) = 0.14,p = 0.71), the temperature values used for the theoretical modelling were calculated as monthly means between surface and bottom temperature. We used two fractionation equations developed for biogenic carbonate to model *δ*^18^*O_Otolith_*. The first equation was proposed by Patterson et al. (1993) as a general equation for freshwater fish, the second equation was proposed by Geffen (2012) for a marine species (plaice, *Pleuronectes platessa* Linnaeus, 1758). These two equations were chosen because no species-specific fractionation equation exists for pike, and Baltic coastal lagoons cannot be easily classified as fully marine or fully freshwater. The two equations took the form

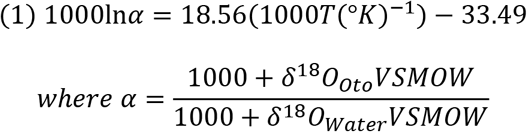

and

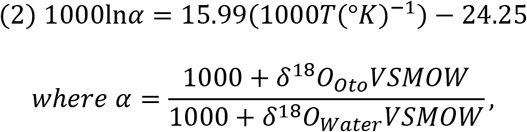

where *T* is the temperature in Kelvin (K), *α* is the fractionation factor between *δ*^18^*O_Water_* and *δ*^18^*O_Otolith_*. Linear mixed effects models were calculated using the lme4 package version 1.1.26 (Bates et al. 2015) to test for the effects of water temperature and *δ^18^O_Water_* on theoretical *δ*^18^*O_Otolith_*, with water temperature and *δ*^18^*O_Water_* as fixed effects and month of measurement as random effect. By keeping one of the two parameters constant while varying the other parameter by its own standard deviation, we assessed the impact of the variability of these two parameters on the predicted *δ*^18^*O_Otolith_* values. All statistical analyses for this study were performed in the statistical programming language R (version 4.05, R core development team 2021) using RStudio desktop version 1.4.1106.

### 2.7. Aging procedure

Aging in this study was performed on high resolution images using the OtoJ macro (Norbert Vischer, Côme Denechaud, Anders Thorsen) within the plugin ObjectJ of imageJ. Image names were randomized prior to aging. The aging procedure consisted of two separate steps performed by three different age readers, named reader 1, 2 and 3 in the following. In a first step, age estimates for the comparison of hard structures were produced for otoliths and scales by readers 1 and 2, where reader 1 was an experienced reader with more than 2000 hard structures read over a time span of more than two years, and reader 2 was inexperienced (no fish hard structures read prior to this work). For scale age estimates, both readers estimated three scales per fish, and the mean age of the three estimates was calculated to reduce bias introduced by individual scales with poor readability. Otoliths were estimated once by both readers. In a second step, to obtain a corroborated age estimate enabling accuracy and bias estimation, we performed a one-to-one mapping of *δ*^18^*O_otolith_* profiles to otolith pictures, aligning the plotted profile to the visible sampling points on the otolith. Precision of the mapping was generally better than 10 *μm*. The images produced in this way were independently assessed by reader 1 and reader 3, both readers with expertise in aging otoliths as well as in interpreting *δ*^18^*O_Otolith_*-profiles on fish otoliths. Because reader 1 also performed step one of the aging, this reader estimated corroborated age only after aging scales and otoliths visually, to avoid introducing bias via prior knowledge from *δ*^18^*O_Otolith_* profiles. The one-to-one mapping approach also allowed an assessment of whether the isotope measurements transect was covering the whole width of the otolith along an axis crossing all annuli. If transects were performed in a way that both reader 1 and reader 3 deemed it possible to have missed annuli, the sample was not used for further analysis. This was the case for two otoliths. In addition to this two-reader corroboration age, an automated peak finding algorithm was defined for the *δ*^18^*O_Otolith_* profiles using the ggpmisc package (Aphalo, 2021). The algorithm was set to detect peaks differing from the mean with at least 50% of the observed isotopic range of an individual and were larger than at least four neighboring values.

### 2.8. Aging accuracy and precision of otoliths and scales

We used the Fisheries Stock Assessment (FSA) package version 0.9.1 (Ogle 2021) to compute basic accuracy and aging bias metrics. To obtain a corroborated age, percent agreement, average standard deviation (ASD), average coefficient of variation (ACV), average absolute deviation (AAD) and average percent error (APE) were calculated and assessed for significant differences and directional bias between the age estimates for the one-to-one mapped images of otoliths and *δ*^18^*O_Otolith_*-profiles provided by reader 1 and reader 3, and for the output of the automated age count algorithm. The automated age count was decided on as the least biased estimate and used for subsequent comparisons between structures and readers. The same metrics as mentioned before were calculated for the otolith and scale age estimates of reader 1 and reader 2, using the corroborated age estimate as reference age. The FSA package tested for a significant difference between the mean corroborated age and the mean-structure-derived age estimate with one-sample t-tests adjusted for multiple comparisons at α=0.05. Wilcoxons signed rank test with a continuity correction (paired by structure) was used to compare the mean coefficient of variation (CV) for all age estimates. We calculated age bias plots comparing mean age estimates for each structure to corroborated age to assess whether there was systematic bias related to structure or reader. We used the McNemar, Evans-Hoenig’s and Bowker symmetry tests within FSA package to test each structure for directional bias associated with the different aging structures.

### 2.9. Modelling population-level growth from size-at-age data

To estimate a population-level growth function for pike, we fitted a von Bertalanffy growth model (Von Bertalanffy, 1938) calculating the size at a given age *L_a_* from the parameters mean maximum adult length *L*_∞_(cm), growth completion coefficient *k* and age at which size is 0 *t*_0_. The function took the form *L*_∞_= *L*_∞_(1 – *e*(^-*k*(*a-t*_0_)^)). Model parameters were estimated in a Bayesian approach, where posterior distributions of the model parameters were determined from a prior distribution and a likelihood distribution. Hamiltonian Monte Carlo (NUTS) sampling was used to sample the posterior distribution of the model parameters in Stan (version 2.21.0). Results of the fitting were processed in R using the rstan package, version 2.21.2 (Stan development team 2020). To select priors, the model was run multiple times on the corroborated age data, with different inputs for the prior distributions of the parameters *L*_∞_ and *t*_0_, following suggestions by Smart and Grammer (2021). We compared the model performances using Leave One Out Information Criterion (LOOIC). The model with the highest LOOIC was used for subsequent growth models. A detailed description of the model comparison along with input parameters for the sampling, diagnostic plots and LOOIC values is provided in the Appendix (section A-B, Figure S1 - S6). The final model was fit using a normally distributed prior determined by the reported maximum length of pike (Froese and Pauly, 2022) for *L*_∞_ and a normally distributed prior for *t*_0_, determined as mean value out of all credible reported estimates for this parameter (Froese and Pauly, 2022). Similar to Smart and Grammer (2021), the growth completion coefficient *k* was set to a uniform distribution bounded from 0 to a maximum probable value of 1, to produce an uninformative prior and allow the model to adapt *k* in order to produce the necessary curvature. All parameters were bounded at zero in order to prevent the model from proposing unrealistic values for the posterior distribution.

Growth models for the comparison of age estimates from different structures were calculated using the age-at-length data from the different calcified hard structures. We used the mean age estimates of both readers for scale and otolith age estimates, yielding a total of three different growth models, a reference growth model using the automated *δ*^18^*O_otolith_*-peak count as corroborated age, a scale-based growth model and an otolith-based growth model. Diagnostic plots for models of all structures are provided in the Appendix.

### 2.10. Assessing fisheries management reference points and optimal size limits

For assessing the impact of bias in aging on pike stock productivity, reference points and optimal harvest regulations, we used a variant of an age-structured population model presented by Ahrens et al. (2020), including size-dependency in maturation, fecundity and natural mortality, and density-dependent recruitment (Table S2). We assumed a constant instantaneous fishing mortality (F), implying a fixed fishing pressure, and size selectivity of the fishery in capture and retention. Deviating from Ahrens et al. (2020), we omitted density-dependence in growth because we were not interested in density-dependence systematics. Rather our aim was to examine how using different growth parameters fitted from the different hard structures affects equilibrium predictions of fisheries metrics in the population model. The parameters informing the model were calibrated to the same stock in the southern Baltic Sea from which the aging data were collected (Table S3). A general description of the model structure is provided in the Appendix (section F).

To estimate production-based (for simplicity, maximum sustainable yield, MSY-based) reference points, we ran model simulations with different values of F, varied in steps of 0.001 *year*^-1^ between 0.0 and 0.6 *year*^-1^, and from that obtained the fishing mortality maximizing the yield (biomass harvested), which was defined as *F_MSY_*. The realized yield and biomass at *F_MSY_* were defined as *MSY* (the maximum sustainable yield) and *B_MSY_* (the biomass at which a fish population is capable of producing *MSY*), respectively. Besides 1.) yield and 2.) biomass which are common objectives for commercial fisheries, our model provided output on 3.) the number of pike being harvested, 4.) the number of individuals being vulnerable to capture and 5.) the number of trophy pike (defined at total length above *L_trophy_*= 100 cm) in the population (Table S2, Eq. 20-24). Metrics 3-5 are more important to recreational fisheries (Ahrens et al., 2020). All five model outputs could be seen as management performance metrics relevant for different stakeholder groups being active in the brackish lagoons (for fishers: 1. and 3., anglers: 3.-5., and conservation: 2.) and were combined as relative values in a log-utility function following Ahrens et al. (2020) (Table S2, Eq. 25). The used log-utility function ensured that each metric was similarly important for looking at suitable compromises in a mixed fishery, where both commercial and recreational fisheries operate (Ahrens et al., 2020). Each model simulation ran for 1000 years and all model outputs were temporally averaged for the last 200 years.

The current minimum-length limit for pike in the Rügen lagoons of 50 cm was used as default for assessing fisheries reference points. To mimic impacts of the growth models on future policies, we varied the minimum-length limit as a common harvest regulation measure between 50 and 100 cm in steps of 1 cm among model simulations with F fixed at 0.2 *year*^-1^, representing the current fishing pressure estimated from stock assessment (Gemert et al., 2022). The one minimum-length limit maximizing the log-utility function composed of the five objectives mentioned above was defined as optimal and could be interpreted as the best compromise among multiple management objectives as in Ahrens et al. (2020).

We obtained the fisheries reference points and the optimal minimum-length limit for three different settings of growth parameterization, based on 1.) scales, 2.) otoliths and 3.) corroborated age, to compare management implications resulting from these three aging methods. We used a pooled approach rather than sex-specific growth functions because of sample size limitations. To account for uncertainty in growth parameter estimation, we randomly sampled *L*_∞_, *k* and *t*_0_ from a normal distribution with the mean and standard deviation derived from measurements of the respective aging method. The normal distributions were truncated to the corresponding 90%-credibility intervals obtained from the data (**2**). For each aging method, 100 random samples of the three growth parameters were taken. For each growth parameter sample, we derived *F_MSY_, MSY* and *B_MSY_* from the simulations with varied F, and the optimal minimum-length limit from the simulations with varied minimum-length limit. Finally, we compared the range in the values of the fisheries reference points and the optimal minimum-length limit among the three aging methods.

### 2.11. Data availability statement

The code used for our analysis is provided in the appendix for reproducibility, the data supporting our findings are available in a public repository on GitHub (https://github.com/Traveller-2909/Pike-age-validation).

## 3. Results

### 3.1. Expected *δ*^18^*O* values in otolith aragonite

Both *δ*^18^*O*_*Water*_ values and water temperature were significant predictors of *δ*^18^*O*_*otalith*_-values (Table S1), however, the variance in *δ*^18^*o_otalith_*-signal induced by temperature observed over a year outweighed the variance which might be introduced by changes in the *δ*^18^*O_Water_* values (Figure **2**). A detailed summary of the model results is provided in the Appendix (Table S1). Observed values from pike otoliths show a comparable amplitude in VPDB values, with maximum values aligning closely with translucent zones and minimum values aligning with opaque zones on the otolith (Figure **3**).

**Figure 2:**
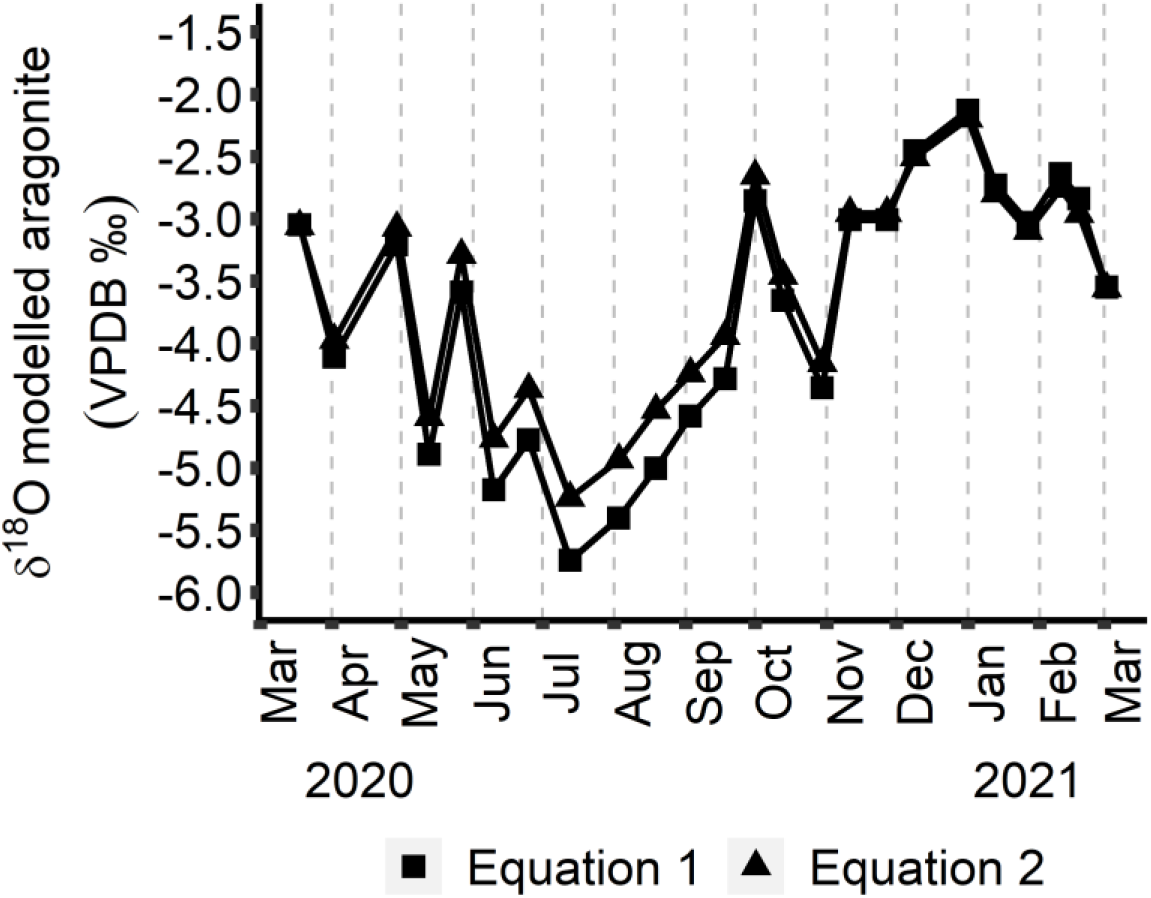
Modelled otolith aragonite values for a hypothetical pike that lived in the Westrügen Bodden chain (WRB) near the water isotope sampling station at Kloster Harbour from March 2020 to March 2021. Predicted aragonite values were calculated from a time-series of biweekly oxygen isotope measurements in the WRB and monthly mean temperatures (surface and bottom). Between April and May 2020 ice coverage at the sampling point of Aichner et al. (2021) prevented sampling for two weeks.

**Figure 3:**
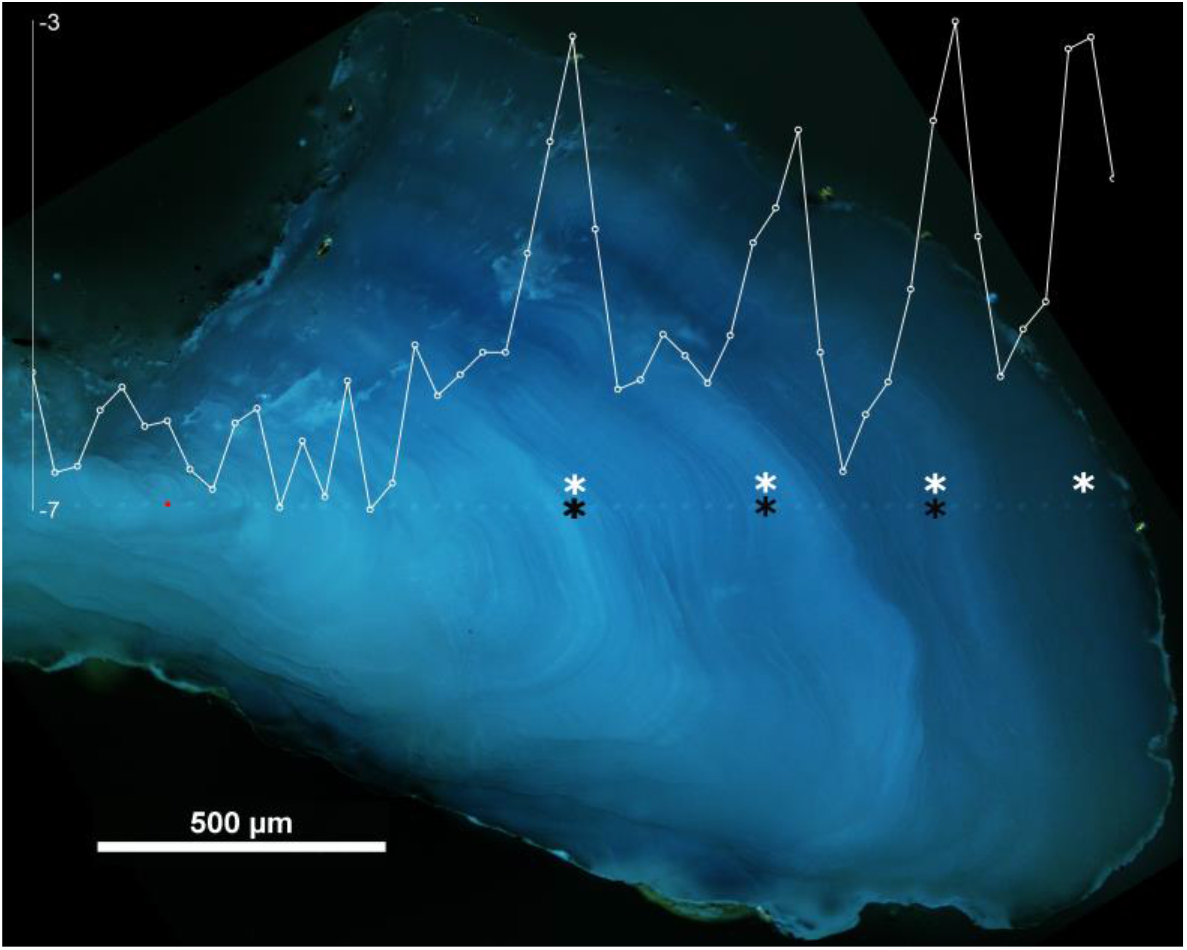
Determined oxygen isotope values in a northern pike otolith caught in Summer 2020 in WRB. The oxygen isotope profile is plotted onto the visible measurement craters on the otolith (dark field image) with a precision of <10 *μm*, isotope values shown on the y-axis are scaled relative to VPDB. Black asterisks mark the aging sequence performed by visual age estimation on the otolith without the isotope profile. White asterisks mark the age estimate performed in combination with isotope profiles.

### 3.2. Obtaining a corroborated age estimation

The age readings of reader 1 and reader 3 on the otolith images combined with *δ*^18^*O_otolith_*-profiles were not unambiguous (between-reader agreement: 36.3%, average coefficient of variation (ACV): 14.8%). Symmetry tests revealed directional between-reader bias (McNemar *χ*^2^= 15.5 with 1 df, p < 0.001; EvansHoenig *χ*^2^= 18.4 with 4 df, p < 0.05). Mean age estimates differed between reader 1 and 3 for the age of 5 years (One sample t-test, t = −4.3, adj. p < 0.05) and 8 years (One sample t-test, t =-4.2, adj. p < 0.05). While there was evidence for directional bias between reader 3 and the automated age count (McNemar *χ*^2^= 7.5 with 1 df, p < 0.05; EvansHoenig *χ*^2^= 11.5 with 4 df, p < 0.05), we found no directional bias between reader 1 and the automated age count (McNemar *χ*^2^= 0.3 with 1 df, p = 0.59; EvansHoenig *χ*^2^= 3.6 with 4 df, p = 0.46). There were no significant differences in mean age estimates between both readers and the automated age count over all age classes (one-sample t-tests, significance level *α* = 0.05). Comparing age estimates in Figure 4 between automated age count and both reader 1 and 3 age estimates, the automated age count lies in between the two readers for most age classes, with a slight underestimation of fish > 10 years compared to the two readers, but this was not significant (one-sample t-tests, significance level *α* = 0.05). In light of these findings, the automated age count was used as corroborated reference age estimate for subsequent analysis.

**Figure 4:**
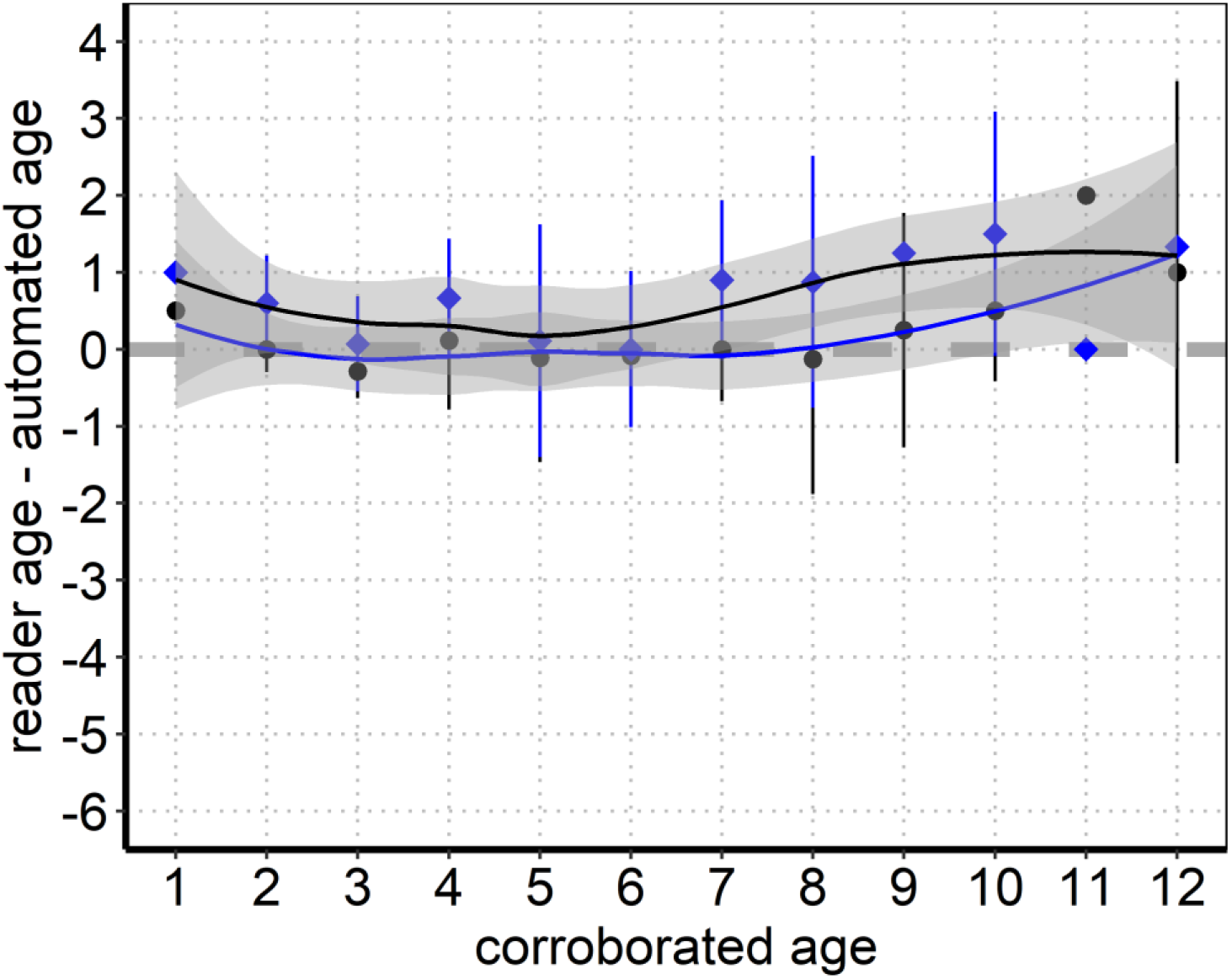
Age-biasplot of corroborated age estimates between two readers and automated peak detection algorithm based on *//delta*^18^*O_Otolith_*-profiles. Automated peak detection age was used as reference age, black dots and line with grey confidence interval show the difference between corresponding age estimates of reader 1 and the reference, blue dots and line show the difference between corresponding age estimates of reader 3 and the reference. Confidence intervals are shaded in grey

### 3.3. Aging accuracy and bias of otoliths and scales

Agreement between age estimates derived from otoliths and scales with the corroborated reference age was lowest for the scale age estimates (27.5% and 23.1% for reader 1 and 2 respectively, Table **1**). Agreement was best for otolith age estimates (36.3% and 35.2% for reader 1 and 2 respectively,**1**). Scale age estimates showed an overall larger bias, as variation coefficients were significantly higher for scales than for otoliths (Wilcoxons signed rank test, reader 1: W = 3394.5, P < 0.05; reader 2: W =3285.5, P < 0.05). We found evidence of aging bias in scale age estimates for both readers 1 and 2, as reader 1 underaged pike at reference ages 7 and 11 (one-sample t test, age 7: *t*(87) = —4.4, p < 0.05; age 11: *t*(87) = —9.8,p < 0.05), and reader 2 overaged pike at reference age 3 (*t*(87) = 3.92,p < 0.05) and underaged pike at reference age 11 (*t*(87) = —8.66,p < 0.05). We found no evidence for under- or overaging in the age estimates from otoliths in both readers, as indicated by non-significant one-sample t tests (significance level *α* = 0.05). We revealed directional age bias between corroborated age and structure age for scale age estimates in both readers using symmetry tests. Reader 1 generally underaged older pike in the age agreement tables (McNemar *χ*^2^= 10.2 with 1 df, p < 0.01, EvansHoenig *χ*^2^= 15.6 with 5 df, p < 0.01), while Reader 2 overaged young age classes and underaged older age classes (McNemar *χ*^2^= 8.2 with 1 df, p<0.01, EvansHoenig *χ*^2^= 13.7 with 6 df, p<0.05). No evidence for directional bias in otolith age estimates could be detected, as indicated by non-significant symmetry tests (significance level *α* = 0.05).

**Table 1:** Summary of aging precision and accuracy metrics.

| Structure | N | % agreement | $\pm 1$ year | $\pm 2$ years | Average coefficient of variation [%] | Average absolute deviation | Average percent error [%] |
| --- | --- | --- | --- | --- | --- | --- | --- |
| Reader 1 scale age | 91 | 27.5 % | 61.5 % | 83.5 % | 24.5% | 0.7 | 17.3% |
| Reader 2 scale age | 91 | 23.1 % | 47.3 % | 73.6 % | 30.0% | 0.8 | 21.2% |
| Reader1 otolith age | 91 | 39.6 % | 78 % | 94.5 % | 16.1% | 0.5 | 11.4% |
| Reader 2 otolith age | 91 | 35.2 % | 74.7 % | 91.2 % | 18.2% | 0.5 | 12.9% |

Comparison of age readings from different structures reveal a marked decline of scale age estimates from the 100 % agreement line for reader 1 in older age classes, starting at age class 7 (figure **5** C). For reader 2, scale age estimates deviate from the 100 % agreement line both for ages younger than 6 as well as for ages older than 9, with the effect being stronger at age 11 (Figure **5** D). Confidence intervals for both otolith estimates largely overlapped with the 100% agreement line for all ages estimated, and no significant deviations of mean structure age from reference age were detected. (Figure **5** A - B).

**Figure 5:**
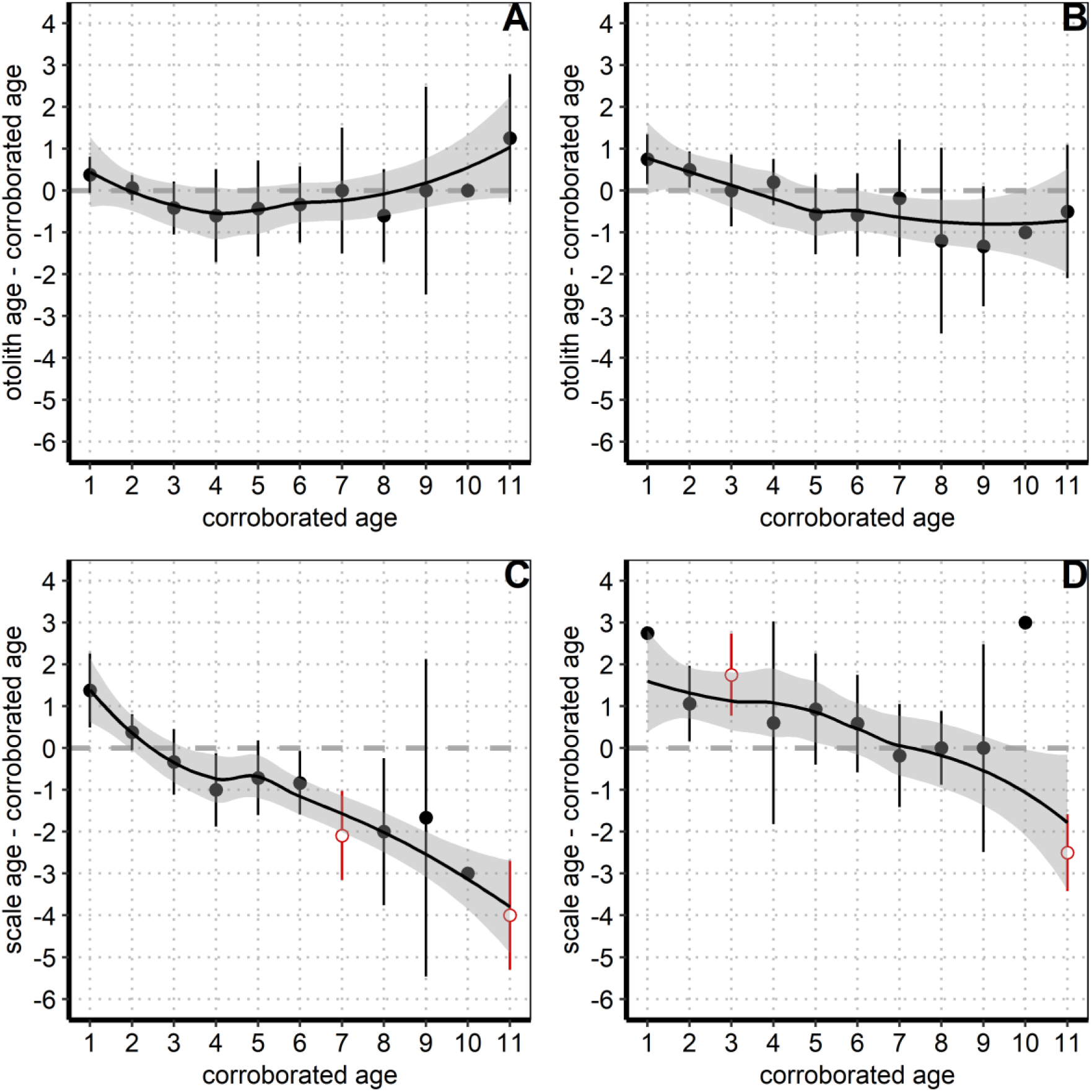
Age bias plots showing the difference between structure age and corroborated age plotted against corroborated reference age. Negative values indicate underaging, positive values indicate overaging by the respective structure. Red points indicate a significant result of one-sample t-test for differences between mean structure age and corroborated reference age. A) Reader 1 otolith age estimate; B) Reader 2 otolith age estimate; C) Reader 1 scale age estimate; D) Reader 2 scale age estimate

### 3.4. Population-level growth of pike using age data from different aging structures

The estimates of population-level von Bertalanffy growth curves did not differ significantly between otolith and corroborated size-at-age data, as indicated by overlapping outer 90% and inner 80% credibility intervals (Figure **6** A-B). The growth curve based on scale-derived size-at-age data differed from the corroborated curve before and after age 7, as shown by non-overlapping 80% inner credible intervals, however, outer 90% credible intervals overlapped slightly between the scale-based and corroborated curves (Figure **6** C). The scale-derived growth curve predicted smaller size-at-age for ages younger than 7 and larger size at-age for ages older than 7 (Figure **6** C), with the effect being stronger for older pike. Scale-aged pike thus show a larger terminal length and lower juvenile growth as indexed by *k*.

**Figure 6:**
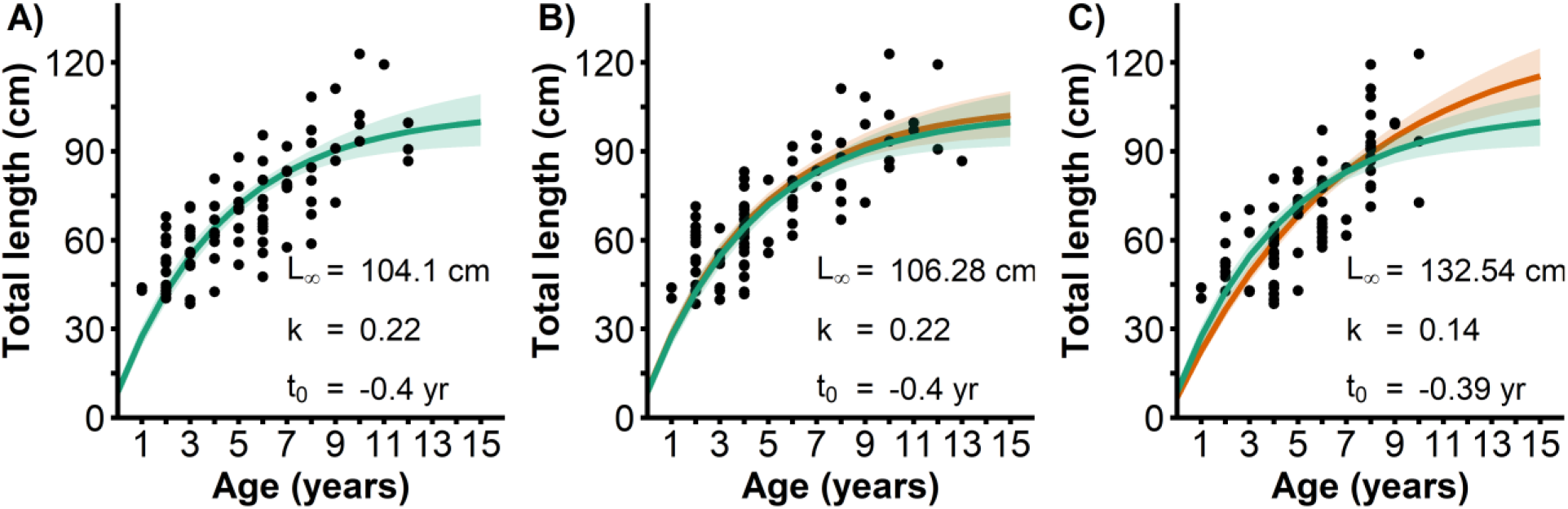
Von Bertalanffy growth curves for all structures and reader age estimates. For reference, the growth curve for corroborated age shown in A) was plotted in green color alongside the curves for different structures in orange color. A) Growth curve for corroborated age estimates; B) Growth curve for otolith age estimates; C) Growth curve for scale age estimates

Growth parameters pooled across sexes that were derived from corroborated age ranged from 91.7 cm to 116.6 cm for *L*_∞_, from 0.16 to 0.28 for *k* and from −0.55 to −0.25 for *t*_0_ based on 90% outer credibility intervals. Growth parameters calculated from otoliths were similar to the parameters calculated using corroborated pike age, ranging from 95.5 cm to 117.2 cm for *L*_∞_, from from 0.17 to 0.27 for *k* and from −0.55 to −0.25 for *t*_0_ (Table **2**). Scale-informed estimates for *L*_∞_ were higher than for the corroborated ages, ranging from 111.1 cm to 153.9 cm, while estimates for *k* were lower, ranging from 0.1 to 0.18, based on 90% outer credibility intervals (Table **2**). Estimates for *t*_0_ were similar between all structures (Table **2**). There was higher uncertainty in parameter estimates for *L*_∞_ and *k* based on scale size-at age data, as the the credible intervals of the posterior distributions for the scale age estimates were wider than those of corroborated age and otoliths. There was evidence for differences in posterior distributions between scale-informed and corroborated parameter estimates, as the inner 80% credibility intervals did not overlap between scale-informed parameters and corroborated parameters. However, there remains some uncertainty about the magnitude of the effect, as outer 90% credibility intervals overlapped for the parameters estimated. MCMC area plots showing the credibility intervals of posterior distributions for all parameters are provided in the Appendix (Figure S13-S16).

**Table 2:** Summary of von Bertalanffy growth function parameters from different aging structures. Outer 90% credibility intervals of the parameter estimates are given in brackets.

| Sample | N | $L_{\infty}$ [cm] | Body growth coefficient k | Age at size 0 [yr] |
| --- | --- | --- | --- | --- |
| Corroborated age | 86 | $104.1 \pm 12.45$ | $0.22 \pm 0.06$ | $-0.4 \pm 0.15$ |
| Otolith age estimate | 86 | $106.3 \pm 10.85$ | $0.22 \pm 0.05$ | $-0.4 \pm 0.15$ |
| Scale age estimate | 86 | $132.5 \pm 21.4$ | $0.14 \pm 0.04$ | $-0.4 \pm 0.15$ |

### 3.5. Fisheries management reference points and optimal minimum-length limit

The fishing-induced mortality at which maximum sustainable yield is achieved (*F_MSY_*), the maximum sustainable yield (*MSY*) and the corresponding biomass at MSY (*B_MSY_*) were found to be substantially lower for model simulations with a growth curve parameterized with scale-based size-at-age data compared to those parameterized with data derived from otoliths or corroborated age (Figure **7** a-c, **8** a,b). For simplicity, we refer to these three model setups as “scale model”, “otolith model” and “corroborated age model” in the following. The otolith model and the corroborated age model yielded a similar level for all three fisheries management reference points/productivity levels. Considering the median values, the scale model underestimated *F_MSY_* compared to the corroborated age model by 12%, MSY by 37% and *B_MSY_* by 27%, while underestimation by the otolith model was marginal (between 1 and 3%) for all three reference points. In contrast to yield and biomass, the amount of large, trophy pike in the population was estimated to be larger with the scale model relative to simulations for the otolith and corroborated age growth models (Figure **8** c), given the higher values of the average asymptotic length *L*_∞_ resulting from age underestimation (Table **2**).

**Figure 7:**
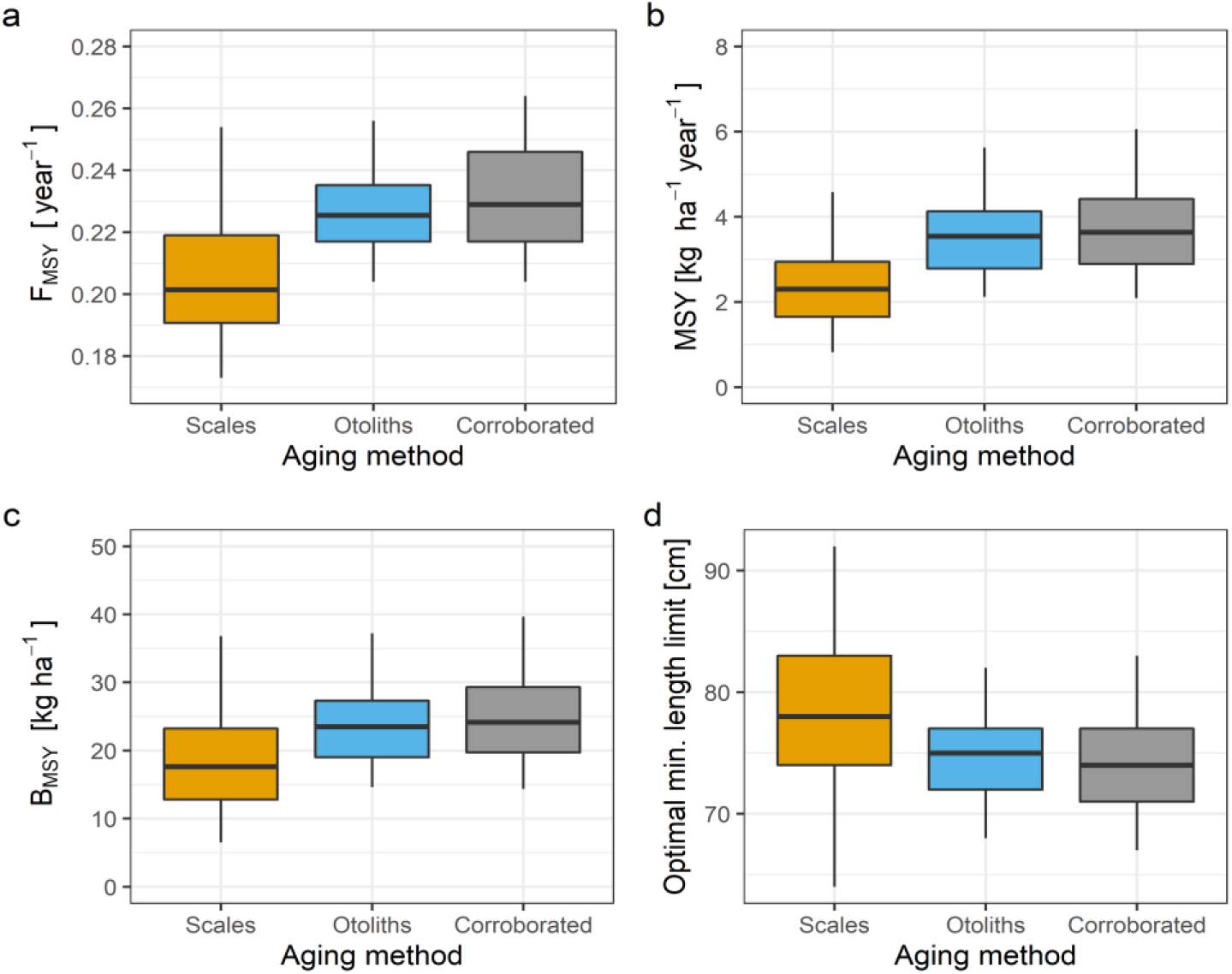
Model results on fisheries reference points *F_MSY_*, MSY and *B_MSY_*, and optimal minimum-length limit, using growth parametrization informed by three different ageing methods. For each ageing method, 100 random growth parameter samples were taken from normal distribution within the 90%-credibility interval of the respective data, accounting for uncertainty in growth estimation. For each sample, the reference points and optimal harvest regulation were obtained from individual model simulations. The variation in the model output shown in the boxplots (outliers not shown) displays the resulting output range in the simulations and allows to compare the implications of the three different ageing methods on fisheries reference points and optimal harvest regulation, under explicit consideration of parameter uncertainty.

**Figure 8:**
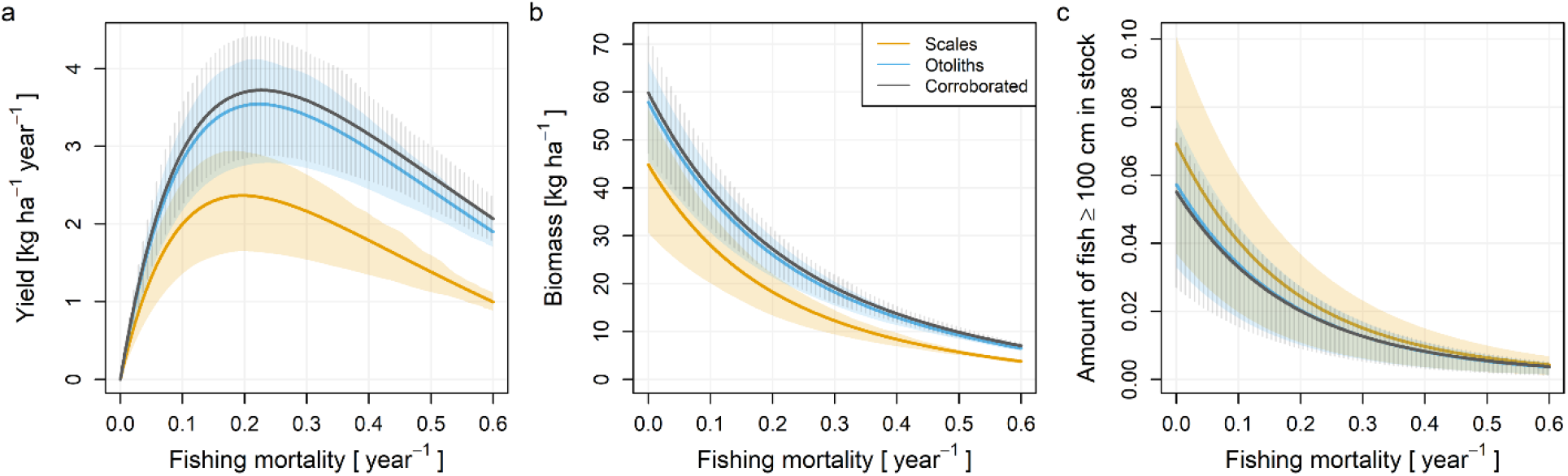
Modelling results on yield, biomass and amount of trophy fish in the population depending on the instantaneous fishing mortality for different growth curve parametrizations based on scales, otoliths and corroborated age. The lines represent mean values of the simulation replicates with different growth parameters, randomly sampled from normal distribution within the 90%-credibility interval estimated from the data of each ageing method. The shaded area equals the respective interquartile range.

The variation in the fisheries reference points among simulation replicates was highest for the scale model (Figure **7** a-c), given the larger relative uncertainty in *L*_∞_ and k estimates (range of 90%-credibility interval relative to mean) based on scales compared to otoliths and corroborated age (Table **2**). Nevertheless, the overlap of the interquartile ranges of the scale model and the corroborated age model (or otolith model) was very low for *F_MSY_* and *MSY*, and low for *B_MSY_* (Figure **7** a-c). This indicated that the difference in the estimated productivity and exploitability between the scale model and the corroborated age model (or otolith model) held also when accounting for parameter uncertainty.

The models with growth curves informed by different aging methods varied also in their management implications for the optimal minimum length limit, but effects were relatively smaller compared to the fisheries reference points (Figure **7** d). The optimal minimum-length limit that maximized the multi-objective utility function was highest for the scale model (78 cm), while it was ~4% lower for the otolith model (75 cm) and ~5% lower for the corroborated age model (74 cm). Variation in the optimal minimum-length limit was relatively large for the scale model (Figure **7** d), given high relative uncertainty in *L*_∞_ estimated based on scales (Table **2**). We tested also for harvest slots as another type of size limit where a maximum size limit is added to the current minimum size limit, but we found no differences in the optimal maximum size limit between the scale and otolith/corroborated age models (Figure S20, 21).

## 4. Discussion

Otoliths produced accurate (as judged against oxygen isotope corroborated age as reference) and largely unbiased age estimates for Baltic pike. By contrast and in support of the first hypothesis, scales were found to be a less reliable aging structure with poor accuracy and precision, shown to be systematically biased, overestimating the age of young pike in one reader and markedly underestimating ages from older individuals for both readers, starting at age 7. However, the two readers varied in the degree of underaging of old pike, suggesting there might also be reader effects. Von Bertalanffy growth curves derived from otolith age estimates were similar to the growth curves produced from corroborated age estimates. Bias introduced by scale underaging of old pike and overall lower accuracy of scale age estimates resulted in a strong effect on growth parameter estimates, in particular an overestimation of the mean terminal length *L*_∞_, and underestimation of the mean metabolic parameter *k* in the von Bertalanffy growth equation relative to corroborated age estimates. This result supported the second hypothesis, i.e., varying age estimates derived from different aging structures had a relevant effect on estimates of population-level von Bertalanffy growth equation parameters. Last, through age-based population models, we found that parameters of growth models using scale-derived size-at-age estimates translated into differing estimates of reference points related to population productivity and also affected management-relevant advice on optimal size-limits, specifically minimum-length limits. This finding is in line with our third hypothesis. Basing fisheries production estimates on scale aging resulted in lower estimates for absolute *MSY, F_MSY_* and *B_MSY_* and slightly higher estimates for optimal minimum length limit and the amount of trophy fish in a population, relative to those inferred from otolith increment (or isotope-corroborated) growth models.

### 4.1. Age corroboration and the value of scales

We based our corroborated ages on coincidence of cyclical variations in *δ*^18^*O* values in the otoliths of pike and annuli. The water temperatures in the shallow lagoon systems were comparable over the whole depth gradient, implying that the majority of the individual fish had experienced the same large seasonal temperature amplitude throughout a year. We modeled the pike otolith aragonite *δ*^18^*O* based on in-water temperature and water *δ*^18^*O* values, demonstrating that the large temperature amplitude outweighed the effects of isotopic variability in the environment on predicted *δ*^18^*O_Otolith_* values. For this study, we used measured values for water *δ*^18^*O* from Aichner et al. (2021) at a sampling site well within our study area to simulate the relative influence of *δ*^18^*O_Water_* on pike *δ*^18^*O_Otolith_*, with the limitation that we simulated this for only one of the three major areas, assuming that the mean yearly variation in *δ*^18^*O_Water_* was constant across the three major lagoon systems. The lagoon this was simulated for exhibits the highest annual salinity fluctuations out of the three study areas (Schlungbaum and Baudler, 2001), which is a strong predictor for *δ*^18^*O_Water_* (Aichner et al., 2021). It is therefore likely that our simulation adequately addressed the seasonal variability of *δ*^18^*O_Water_*, lending robustness to the assumption.

The *δ*^18^*O* values observed in pike otolith SIMS profiles showed a comparable amplitude to the predictions made by our theoretical model. Sources of variations in observed patterns in otoliths are likely attributable to interindividual variation in thermal preference, such as Morissette et al. (2020) reported for lake trout (*Salvelinus namaycush* (Walbaum, 1792)), or environmental fluctuations influencing the isotopic composition of the ambient water, such as evaporative effects (Bowen et al., 2018). Another source for deviations from predicted values could be organic fraction in the aragonite matrix of the otolith. Matta et al. (2013) demonstrated an offset of 1‰ for yellow sole (*Limanda aspera* (Pallas, 1814)) otoliths that were roasted at 450 °C for 1 h to remove organic residues, when compared to untreated otoliths. Matta et al. (2013) found the offset to be consistent along the whole width of the otolith, therefore retaining the overall seasonal temperature signal. We thus assume these effects, should they exist in the lagoons for pike as well, did not blur the patterns we revealed in our work. Also, as we did not aim at deriving absolute temperature values from *δ*^18^*O_Otolith_* values in our study, only the seasonal amplitude of the *δ*^18^*O_Otolith_* values was of interest for corroborating age estimates. In our work, the seasonality of *δ*^18^*O_Otolith_* was shown to outweigh the possibility for lower amplitudes through environmental fluctuations. Hence, we consider the *δ*^18^*O_Otolith_* profiles useful and reliable for inferring the timing of growth checks in pike. This conclusion agrees with Gerdeaux and Dufour (2012), who demonstrated a strong correlation of visible growth checks on pike otoliths with maxima and minima of aragonite *δ*^18^*O* values.

Age estimates based on visual examination of otolith annuli were largely similar to isotope-based age estimates, revealing a high accuracy of this structure for aging pike. Our finding agrees with previous work in freshwater pike by Oele et al. (2015) and Blackwell et al. (2016) demonstrating low bias and high precision of otoliths when aging pike. Scale readings, by contrast, exhibited poor accuracy and precision scores for both the experienced as well as for the inexperienced reader, similar to previous work in pike (Blackwell et al., 2016; Frost and Kipling, 1959; Mann and Beaumont, 1990; Oele et al., 2015). However, the direction of bias differed by reader, rendering it difficult to arrive at clear conclusions as to how exactly scale data are biased. Age bias plots indicated the potential for overaging of young pike and significant underaging of older pike along with a high degree of uncertainty in mean age estimates, but the degree differed strongly between readers. While the more experienced reader 1 only overaged age 1 pike and underaged pike starting at age 7, the less experienced reader 2 overaged over a wide range of young pike sizes, but underaged pike only starting at age 11, suggesting this reader misinterpreted the onset of piscivory or other ontogenetically induced checks and changes in growth rate as annual marks (e.g., Frost and Kipling (1959), Mann and Beaumont (1990), Casselman (1974)). With only two readers for the comparison of scales and otoliths, we are unable to relate our findings to experience effects as we cannot differentiate from experience-independent personal effects in ability to age. However, reader effects are known from other studies in pike (Oele et al., 2015) and other species (Hüssy et al., 2016). We found no impact of reader on symmetry tests, suggesting any systematic bias to be broadly consistent between the two readers in our work. Importantly, the age at which the first significant underaging occurred agreed with the findings of Blackwell et al. (2016) and Oele et al. (2015), who reported significant underaging of pike starting after age 6 using scales. A similar bias as revealed for pike was also reported by Tyszko and Pritt (2017) for largemouth bass *Micropterus salmoides* (Lacepède, 1802). Our results disagree with work on pike by Laine et al. (1991), who reported good accuracy of scales up to an age of 10 years compared to cleithrum age estimates that were validated via a tetracycline marking approach in a lake in Ontario. Relatedly, cleithra and scales were found to reveal consistent results by Pagel et al. (2015), and cleithra were identified before as a suitable aging structure in pike (Casselman, 1974). An explanation for the apparent discrepancy in readability of scales could be related to differences in growth rate in different populations, but also to environmental impacts on structure readability, such as effects induced by the brackish lagoon habitat, similar to what has been reported for Baltic cod (Hüssy et al., 2016). Given the strong correlation of scale length to body length in fish, scales become difficult to read in older fish, as the outer annuli are compressed in old fish, and become indistinguishable (Blackwell et al., 2016; Casselman, 1990; Frost and Kipling, 1959; Mann and Beaumont, 1990; Oele et al., 2015). The critical age after which scale readings become problematic in pike seems vary by reader and will in practice likely also vary by the growth rate of the local population.

In contrast to scales, otoliths continue to grow when fish are in starvation or catabolisis, as otolith growth is linked to the metabolism of the fish rather than to somatic growth (Campana, 1990). While this property complicates back calculations of body size using otoliths (Essington et al., 2022), it may explain the improved readability of this hard structure in many different fish species. Scales, however, have the obvious advantage that they can be sampled non-lethally and are suitable for back-calculation of body size, especially at young ages (Pagel et al., 2015). Thus, the ultimate choice of the aging structures strongly depends on the research objective. More generally, given the systematic impact of scales on underaging of old fish, comparing growth curve parameters with different structures seems problematic as underaging strongly affects estimates of *k* and *L*_∞_. As the minimum, among study comparisons should pay careful attention to the size and age range in the different studies. Further system-specific validation studies with true known age fish are therefore warranted.

### 4.2. Consequences for growth and fisheries reference points

Bias introduced by underaging older fish in scales and overall lower accuracy resulted in overestimation of *L*_∞_ and an underestimation of the growth completion parameter *k*, with parameter estimates for *t*_0_ being largely comparable between all sets of age estimates. These results agree with previous work comparing growth model estimations based on different age estimation methods by Tyszko and Pritt (2017) in largemouth bass. The overestimation of *L*_∞_ by the scale-informed growth models is likely the result of underaging older fish, which lead to an overestimation of size-at-age for older age classes, increasing the steepness of the growth curve, thus leading the function to reach the asymptotic phase at larger sizes. The underestimation of the metabolic parameter *k* for both our study and that of Tyszko and Pritt (2017) is attributable to the inherent negative correlation between *k* and *L*_∞_ parameters (Pilling et al., 2002; Xiao, 1994), and is ultimately caused by the fitting procedure (Pilling et al., 2002; Xiao, 1994). Thus, caution is encouraged when extracting point estimates for parameters derived from Bayesian posterior probability distributions for further analysis. In fact, the estimation of point estimates from biased age data might have consequences when population growth parameters are compared between multiple systems (Quist et al., 2003), or for meta-analysis approaches, where estimates of growth parameters are extracted from multiple studies (Rypel, 2012). For example, a systematically larger *L*_∞_ estimate for scale-read data can bias meta-analyses on fish growth if many studies entered in the meta-analysis used scales for aging as opposed to otoliths. As shown by Tyszko and Pritt (2017) and Campana (2001), use of different structures inducing different levels of bias for age estimation could make comparisons between systems and studies unreliable. Therefore, attention should be paid to which aging structures were used in the respective studies and, ideally, age corroboration or validation studies should be available for the species of interest (Campana, 2001; Tyszko and Pritt, 2017).

Our age-based population model using scale-derived growth parameters underestimated *MSY, F_MSY_* and *B_MSY_* compared to the otolith and corroborated age model, despite higher scale-based estimates for *L*_∞_. This result was attributable to the large influence of the metabolic parameter *k* for productivity in age structured

models (Figure S17), which was underestimated by using scales for aging. A lower *k* implies slower growth rates especially at young ages, raising the age at which fish become vulnerable to a fishery and thus decreases the harvestable stock biomass at a given constant mortality, which is why lower values of *k* (as inferred from scale compared to otolith age estimates) translated into lower sustainable yield (Beamesderfer and North, 1995; Beverton and Holt, 1957). This effect might be even stronger given negative size-dependency in natural mortality, which leads to increased natural mortality of younger age classes at low *k* (Lorenzen, 2008; Lorenzen and Enberg, 2002). The decreased productivity, implied by a low *k*, typically decreases the fishing mortality at which maximum sustainable yield is achieved, i.e., *F_MSY_* (Horbowy and Hommik, 2022), but this effect also depends on the given level of *L*_∞_, see Figure S17.

The reduced *MSY, B_MSY_* and *F_MSY_* predicted by our scale-informed models could motivate fisheries managers to to apply too conservative regulations to limit the fishing mortality. If the fish stock would be managed under a *MSY* objective, an overly conservative approach would lead to a fishing mortality lower than what would generate *MSY*, resulting in lost yield potential at the benefits of reducing the potential for growth overfishing (Hilborn, 2010). By contrast, under the objective of a trophy fishery, which is more oriented towards recreational fisheries (Ahrens et al., 2020), the overestimation of mean adult maximum adult size exhibited by the scale derived ages could lead to biased and overly optimistic expectations towards the enhanced production of trophy fish. This problem was also encountered in the work of Tyszko and Pritt (2017), who found scale-based models to predict a satisfactory amount of trophy fish in largemouth bass populations even at high fishery mortalities, whereas the evidently more reliable otolith-based models estimated a decrease in trophy fish in the population at any given fishery mortality.

In areas with co-exploitation of stocks by commercial and recreational fisheries (as in the lagoon fishery around Rügen), both biomass and catch rate/trophy fish size related objectives are jointly relevant (Gemert et al., 2022), and compromises have to be reached between competing objectives (Ahrens et al., 2020). Using an integrated multi-objective utility function, we found that scale-informed models would favor a higher optimal minimum-length limit (around 80 cm) compared to models informed by otoliths or corroborated age(around 75 cm). Noteworthy, all models suggested optimal minimum-length limits that were larger than the status quo (50 cm). This difference in optimal minimum-length limit was due to the higher amount of large fish predicted based on scales (higher *L*_∞_) that fall into the harvestable category even when increasing the minimum length limit. While the growth models varied in their predictions of the optimal minimum-length limits for multiple objectives, the effects were relatively speaking less than when looking at the productivity-metrics alone. Thus, while we found the scale age growth model strongly affected predictions of MSY-based reference points, the optimal size limit regulations predicted for catch and harvest objectives were rather similar. However, when it comes to quota as a binding harvest regulation, the differences in the recommended quota between the different growth models might be larger given the different productivity estimates for the stock that we revealed.

### 4.4. Limitations

A possible limitation to our approach lies in size-selectivity of sampling methods. Most fisheries methods used to sample fish are selective with respect to size (Frater and Stefansson, 2019; Pardo et al., 2013), which can lead to biased growth estimations (Pardo et al., 2013). In our work, we followed the recommendations of Wilson et al. (2015) and selected our sample from a larger pool of fish caught with different gear. As we compared the output of different sets of age readings among the same sample of fish, varying only the ages as input to the growth models, a systematic error from gear selectivity is unlikely to have affected our results. Using *δ*^18^*O_Otolith_* values to corroborate age readings bears the limitation that temporal resolution of sampling points decreases with increasing age of the fish due to narrower increments. However, resolution in the outermost visible annuli was > 4 SIMS-points for the oldest individuals in our study, which we deemed sufficient for detecting a seasonal temperature signal, in accordance with previous work (Kastelle et al., 2017; Weidman and Millner, 2000). Having only two readers for scales and otolith readings also poses a limitation, as it makes it difficult to distinguish between bias introduced by a given structure from reader-introduced bias. In our age-structured population model, we did not include density-dependent growth, which may cause positive feedback of harvest-induced reduction of biomass of pike growth due to decreased competition and social stress (Edeline et al., 2010). Such feedback will affect the reference points and optimal size limits for the different growth models (Ahrens et al., 2020), but should not qualitatively change our results.

### 4.5. Conclusions and implications

We provide a case study investigating aging bias in fish calcified structures, using high-resolution otolith *δ*^18^*O* chronologies to corroborate annual growth checks in Baltic pike. Our work, first, shows that *δ*^18^*O* data are suitable as a thermal marker in pike, similar to findings in other fish studies (Campana et al., 2020; Geffen, 2012; Morissette et al., 2020). Second, we report age estimates from aging structures with poor accuracy and precision, in our case scales, substantially affect population-level growth and productivity estimates, raising a cautionary note about the influence of aging structures on growth and productivity assessment of fish. Importantly, we found the age bias introduced at the growth model stage to propagate to productivity and yield estimation, possibly leading to suboptimal management suggestions. We can offer three recommendations. First, growth rate comparisons over time in a given area shall ideally be based on the same aging structure and be confined to age and size ranges for which the specific structure is found to be reliable. Second, if the aim is to arrive at an unbiased population level growth assessment or estimates of maximum age, scales are not a suitable aging structure in Baltic pike and alternatives, such as otoliths, should be used. Finally, inclusion of growth models estimated from scales can lead to conservative management when the objective is to maximize biomass yield and to overly optimistic predictions as to the trophy potential of the fishery. The implications of our results further underline the importance of quality control and validation studies when aging fish for ecological inference or to meet fisheries objectives, and to think carefully about which aging structure to use, which will vary depending on study objectives and the inferences that the aging structure is supposed to generate. For the case of Baltic pike, we recommend further validation and corroboration studies with other non-lethally sampled aging structures, such as fin rays.

## 5. Acknowledgements

Funding was provided by the Deutsche Bundesstiftung Umwelt (German Federal Environmental Foundation, DBU, No AZ 20019/634) through a scholarship to TR, by the Leibniz Institute of Freshwater Ecology and Inland Fisheries (IGB) Berlin, and through the European Maritime Fisheries Fund (EMFF) of the EU and the State of Mecklenburg-Vorpommern (Germany) (grant B 730117000069, MV-I.18- LM-004) within the Boddenhecht project. We thank Rob Ahrens for advice on growth modelling and Thomas Mehner and the participants of the writing course run by Thomas Mehner for helpful discussions on the manuscript. We thank two anonymous reviewers for excellent feedback on the manuscript.

