## Supplemental material for "Corroborating age with oxygen isotope profiles in otoliths: consequences for estimation of growth, productivity and management reference points in northern pike (*Esox lucius*)in the southern Baltic Sea"

**Appendix**

**A: Multimodel comparison**

For model 1, we set the mean of the prior distribution for $L_\infty$ to the reported maximum length for pike (150 cm, Caspers et al. (1968)) and the standard deviation to 5% of the mean (7.5 cm),to generate a more restrictive prior. For model 2, the standard deviation of the two priors was set to be less restrictive, to 10 % of the mean. Model 3 was calculated with setting 0 as the mean for the priors for $L_\infty$ and $t_0$, i.e., giving no information on mean maximum length and mean age at which length is 0, only providing a standard deviation covering the range of mean values of the first two parameter sets. The standard deviation was set to 150 cm for $L_\infty$ and to 1 for $t_0$ for this model. All models for this step were run for 10,000 iterations with a burn-in period of 5000 iterations. Models were checked for convergence efficiency and autocorrelation by examining diagnostic plots from the bayesplot R package version 1.8.0 (Gabry 2021). Because no autocorrelation could be detected, no thinning was performed. LOOIC values were calculated for all models using the loo R package version 2.4.1 (Vehtari 2017).

### B: Model diagnostics for multimodel comparison

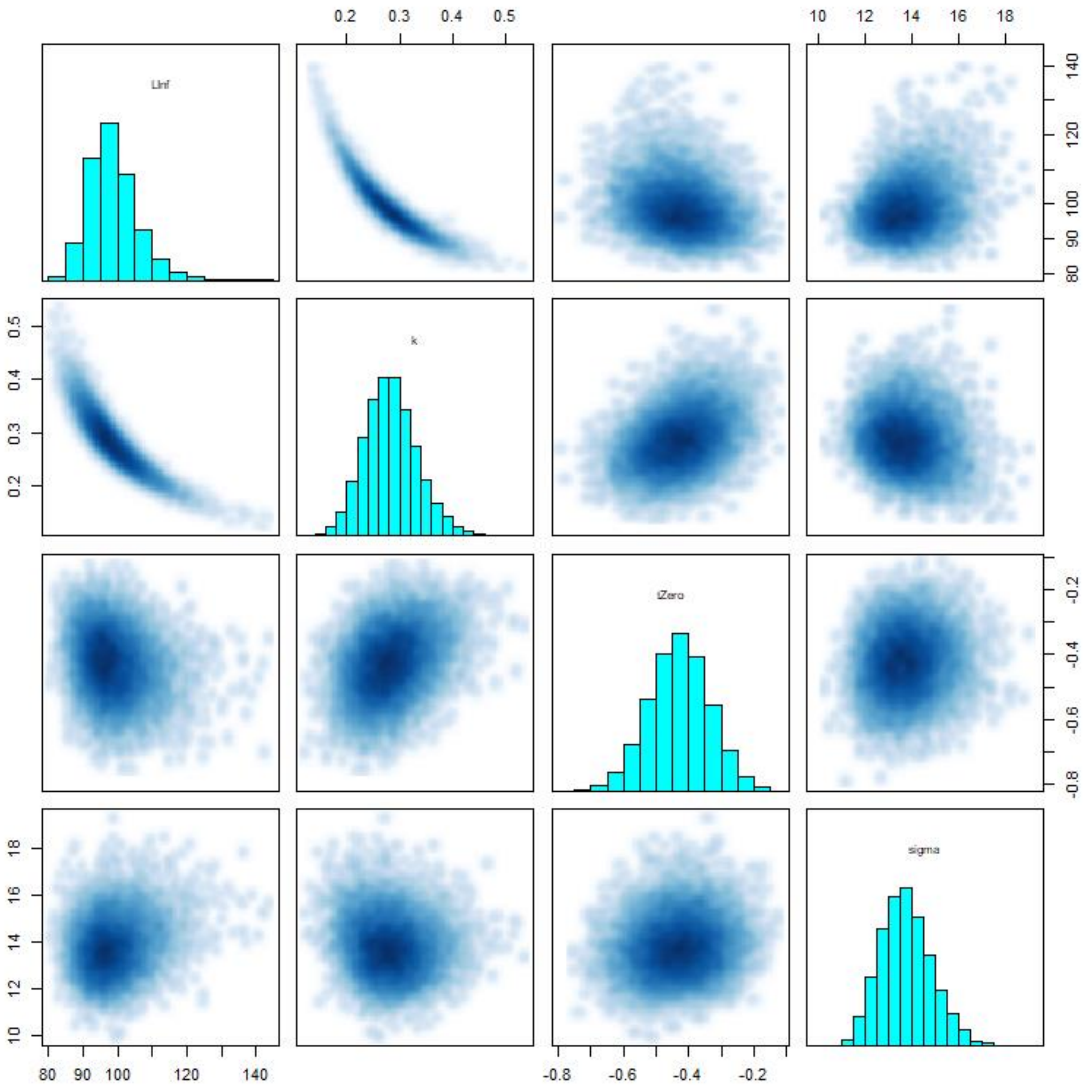


Figure S1: Pairs plot for model 1, showing the interactions between variables used in the model. The inherent negative correlation between $L_\infty$ and *k* is apparent by the presence of multiplicative non-identifiabilities (top left corner). Remaining parameters show even distributions in their respective bivariate plots as well as evenly distributed univariate histograms, indicating no issues of collinearity between variables is present.

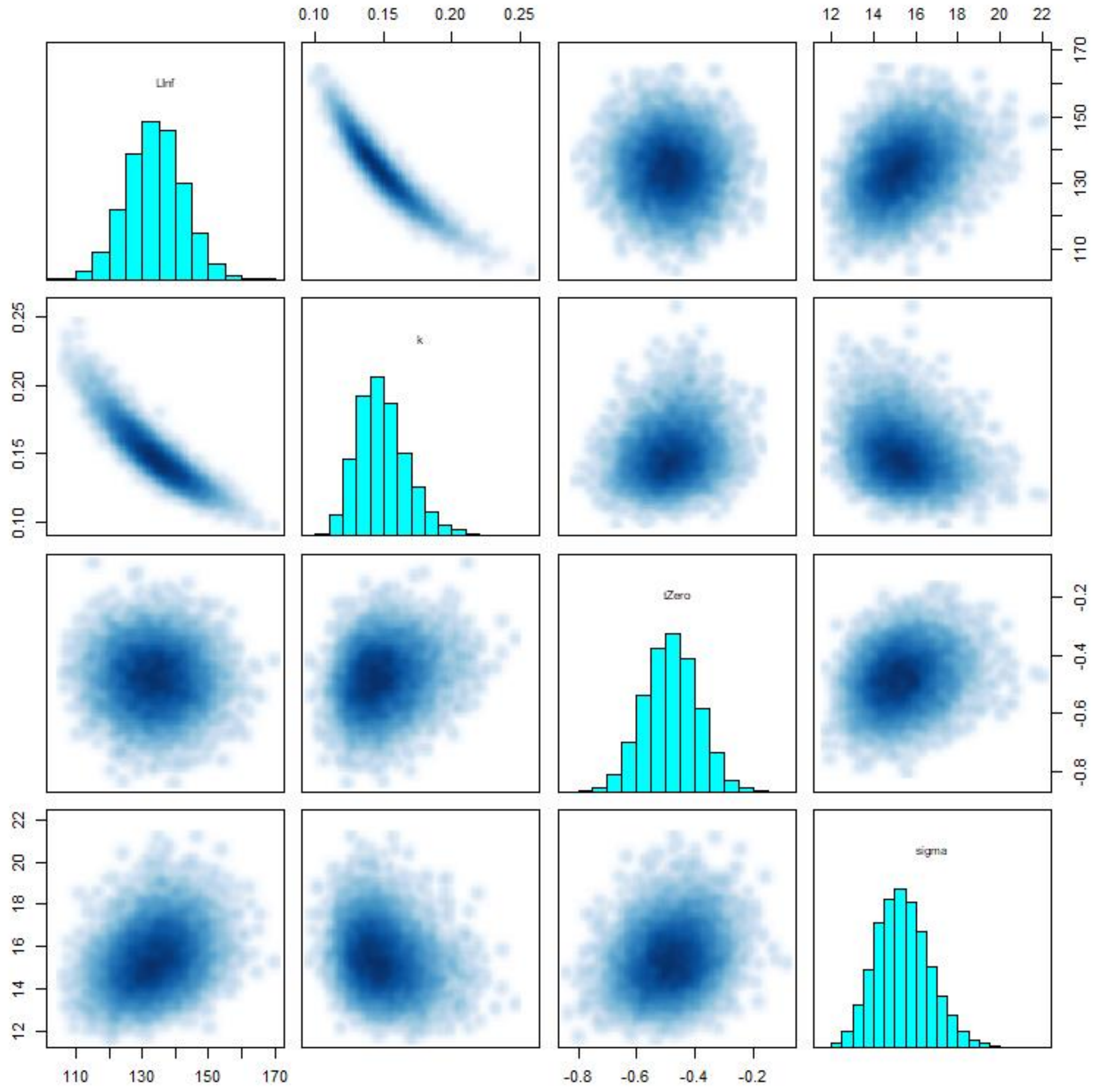


Figure S2: Pairs plot for model 2, showing the interactions between variables used in the model. The inherent negative correlation between $L_\infty$ and *k* is apparent by the presence of multiplicative non-identifiabilities (top left corner). Remaining parameters show even distributions in their respective bivariate plots as well as evenly distributed univariate histograms, indicating no issues of collinearity between variables is present.

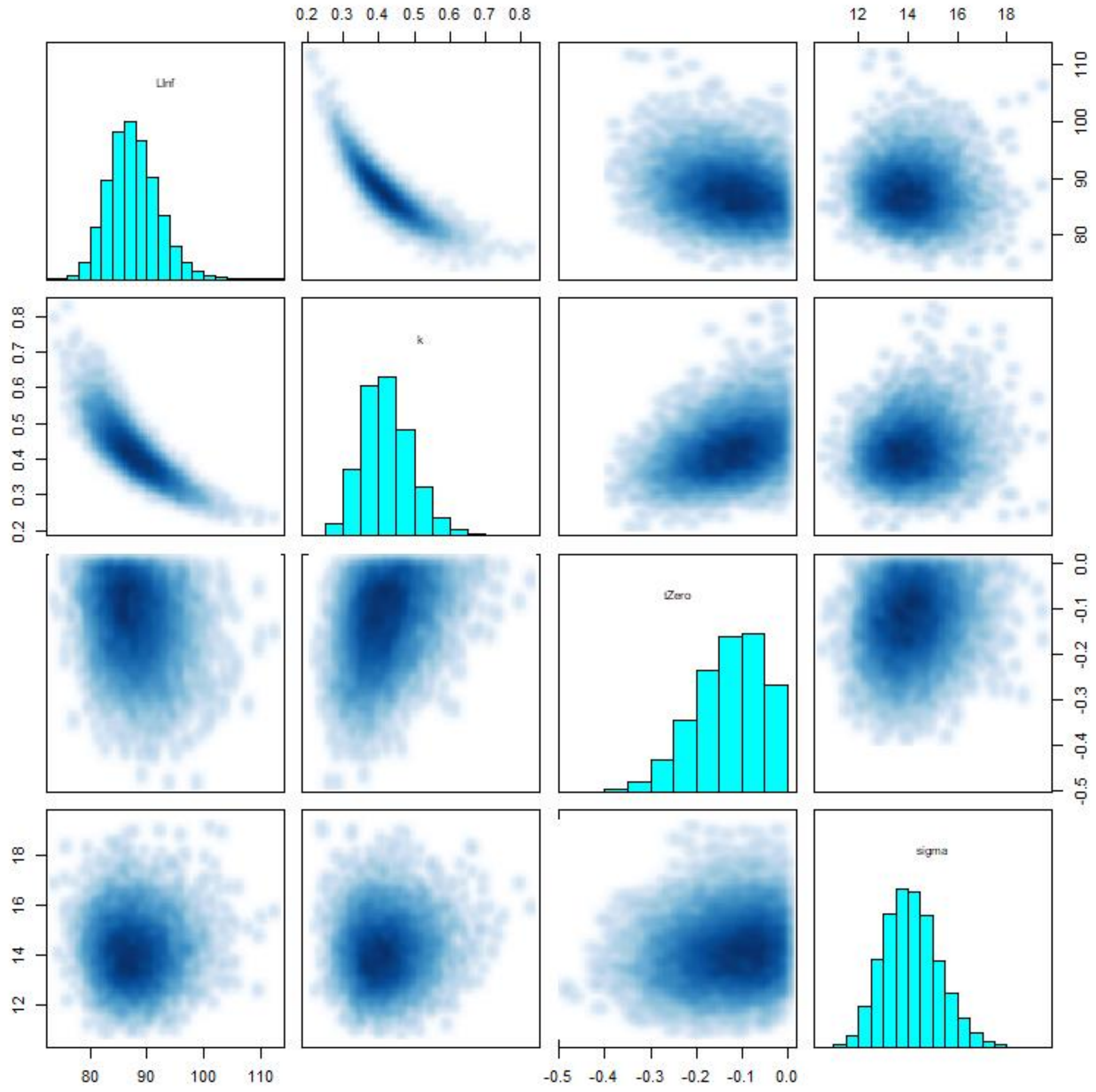


Figure S3: Pairs plot for model 3, showing the interactions between variables used in the model. The inherent negative correlation between $L_\infty$ and *k* is apparent by the presence of multiplicative non-identifiabilities (top left corner). Bivariate plots of k and $t_0$ show a funnel.like hape, indicating sampling problems of the Markov chains. Similar sampling issues are present between $t_0$ and sigma.

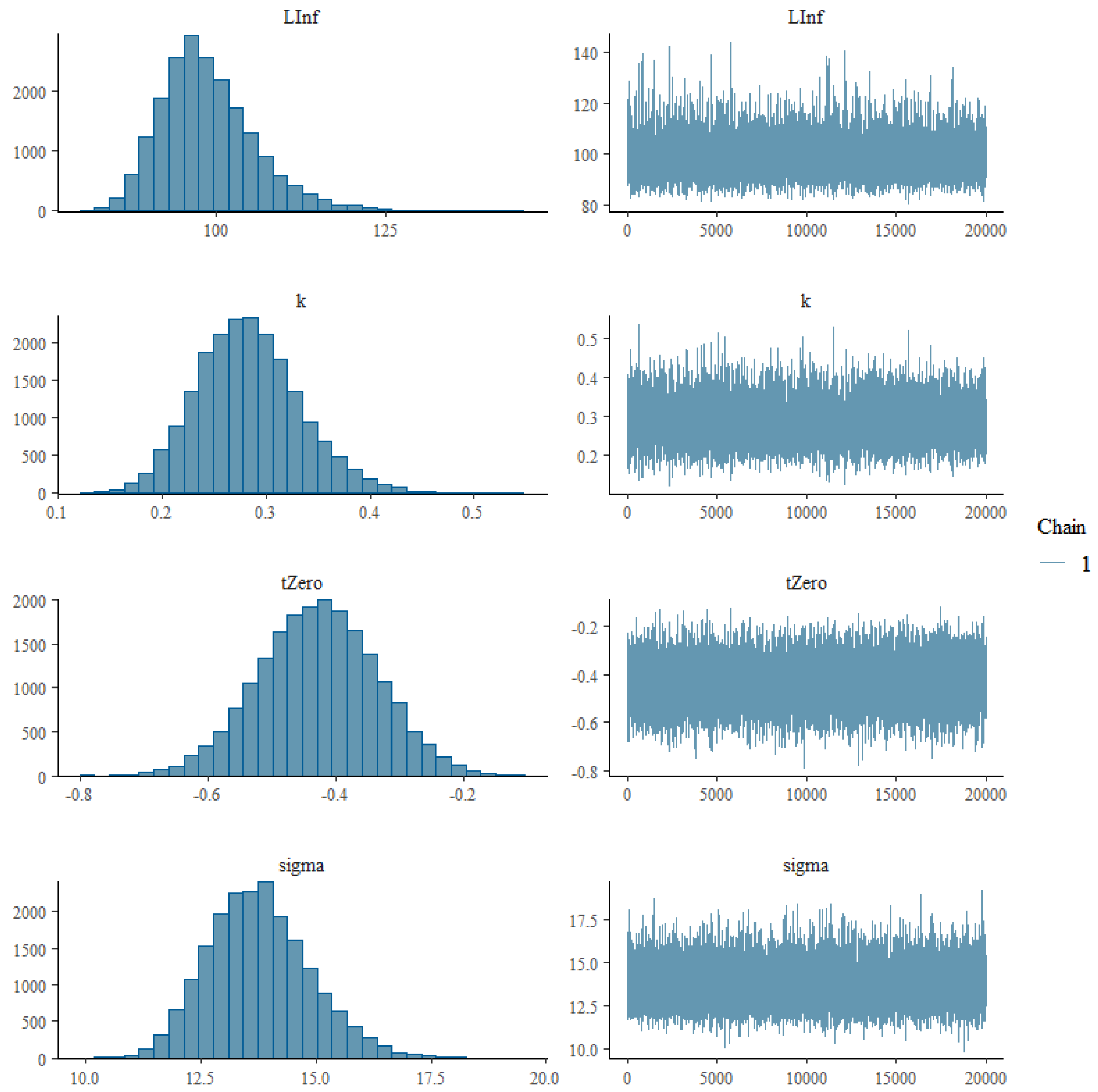


Figure S4: MCMC combo plot for model 1. Traceplots (right side) indicate chains have mixed for all parameters and model efficiency was satisfactory. Parameter histograms (left side) show no problems of multi-modality.

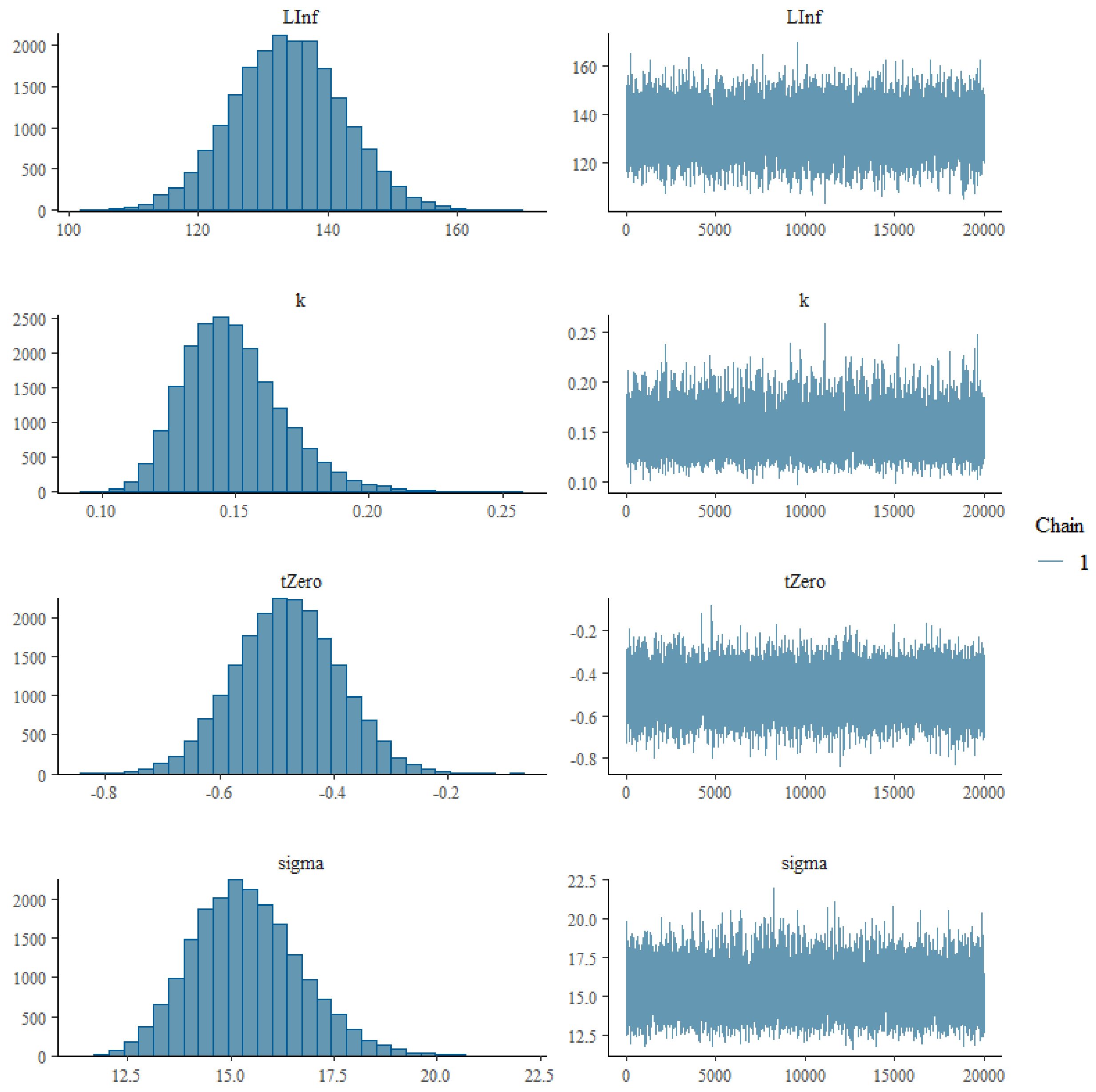

Figure S5: MCMC combo plot for model 2. Traceplots (right side) indicate chains have mixed for all parameters and model efficiency was satisfactory. Parameter histograms (left side) show no problems of multi-modality.

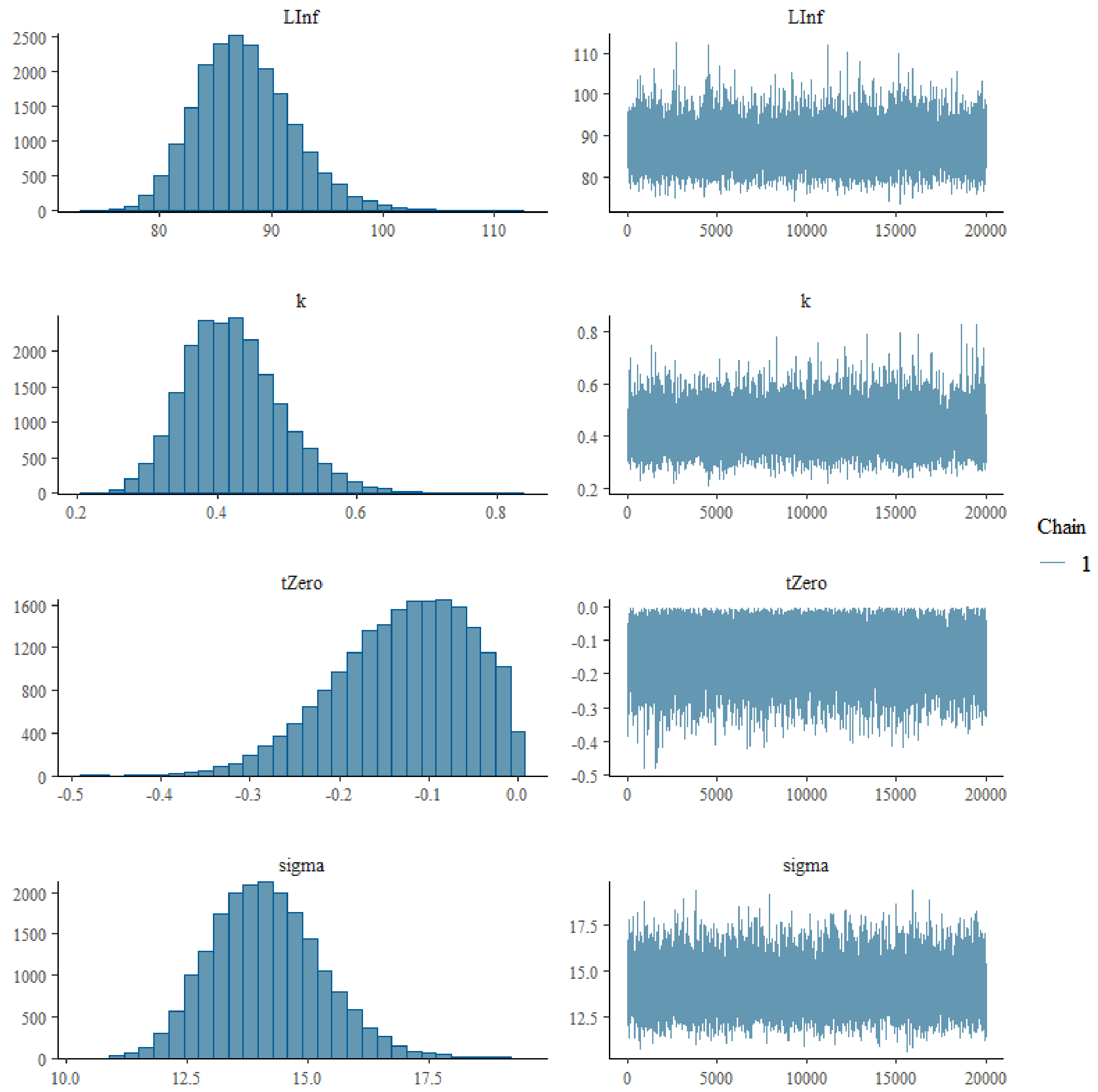


Figure S6: MCMC combo plot for model 3. Traceplots (right side) indicate chains have mixed for all parameters and model efficiency was satisfactory. Parameter histograms (left side) show no problems of multi-modality.

### C: Model diagnostics for structure-specific growth models

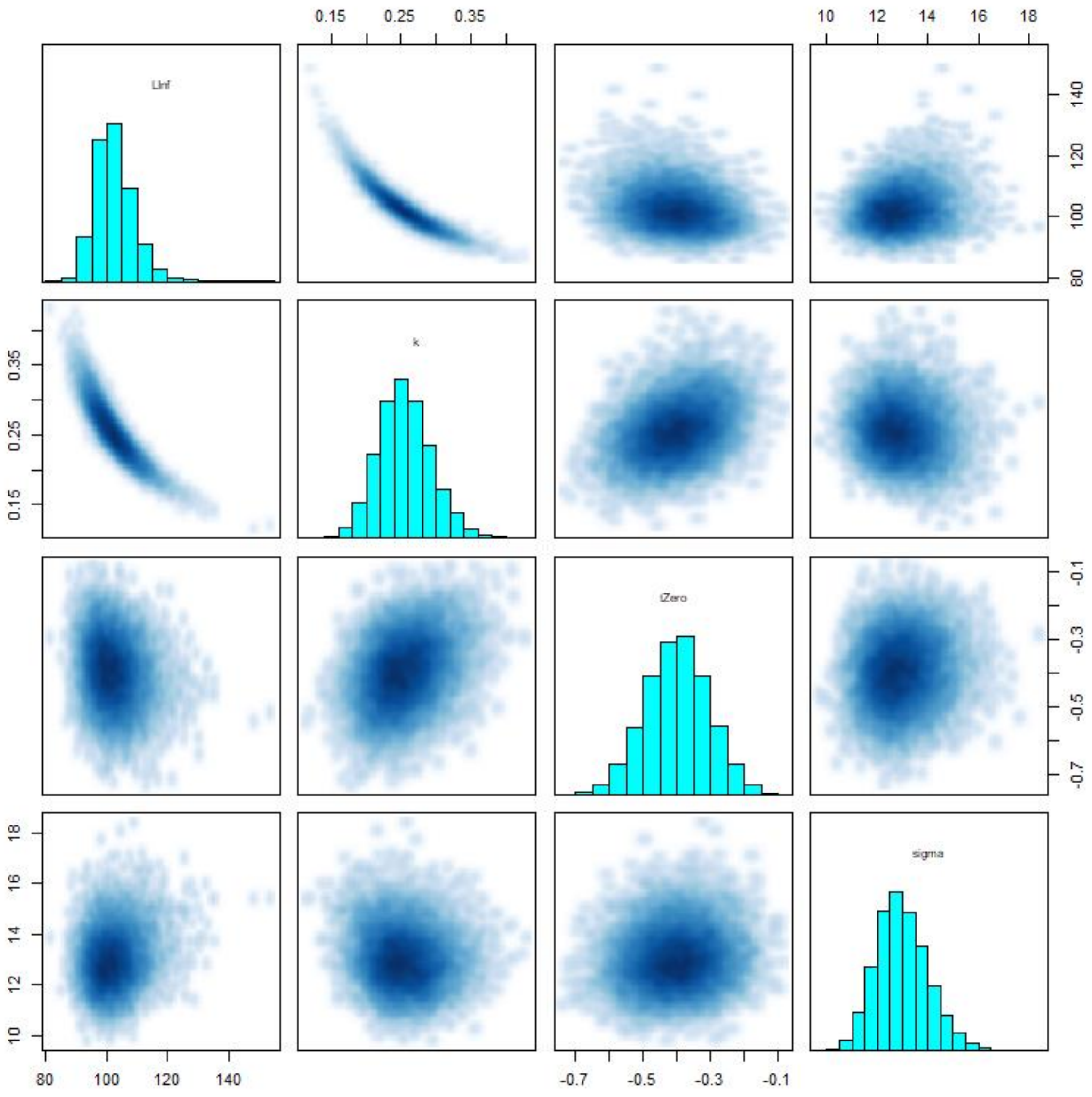


Figure S7: Pairs plot for the corroborated age model, showing the interactions between variables used in the model. The inherent negative correlation between $L_\infty$ and *k* is apparent by the presence of multiplicative non-identifiabilities (top left corner). Remaining parameters show even distributions in their respective bivariate plots as well as normally distributed univariate histograms, indicating no issues of colinearity between variables is present.

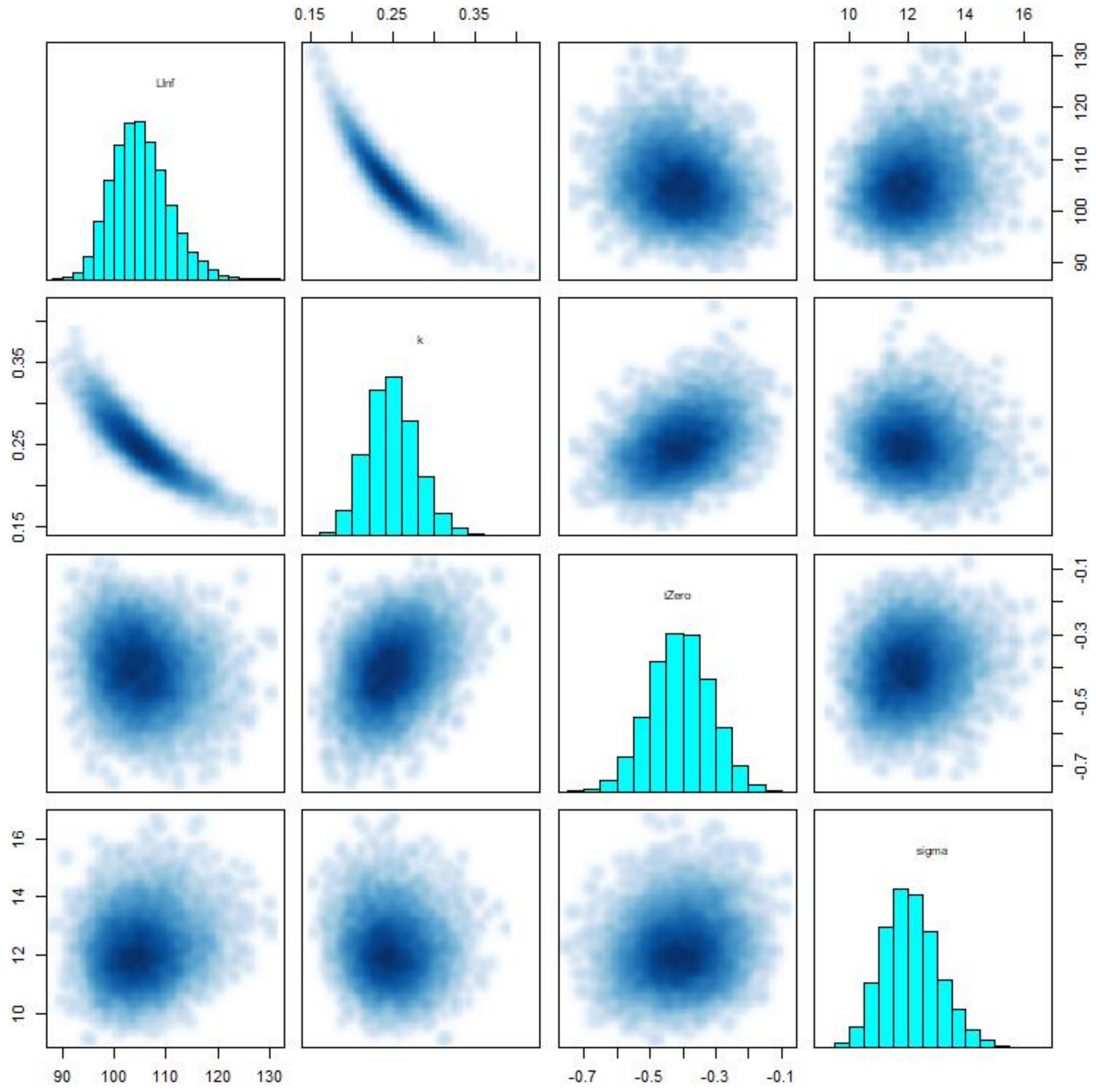


Figure S8: Pairs plot for the otolith age model, showing the interactions between variables used in the model. The inherent negative correlation between $L_\infty$ and *k* is apparent by the presence of multiplicative non-identifiabilities (top left corner). Remaining parameters show even distributions in their respective bivariate plots as well as normally distributed univariate histograms, indicating no issues of colinearity between variables is present.

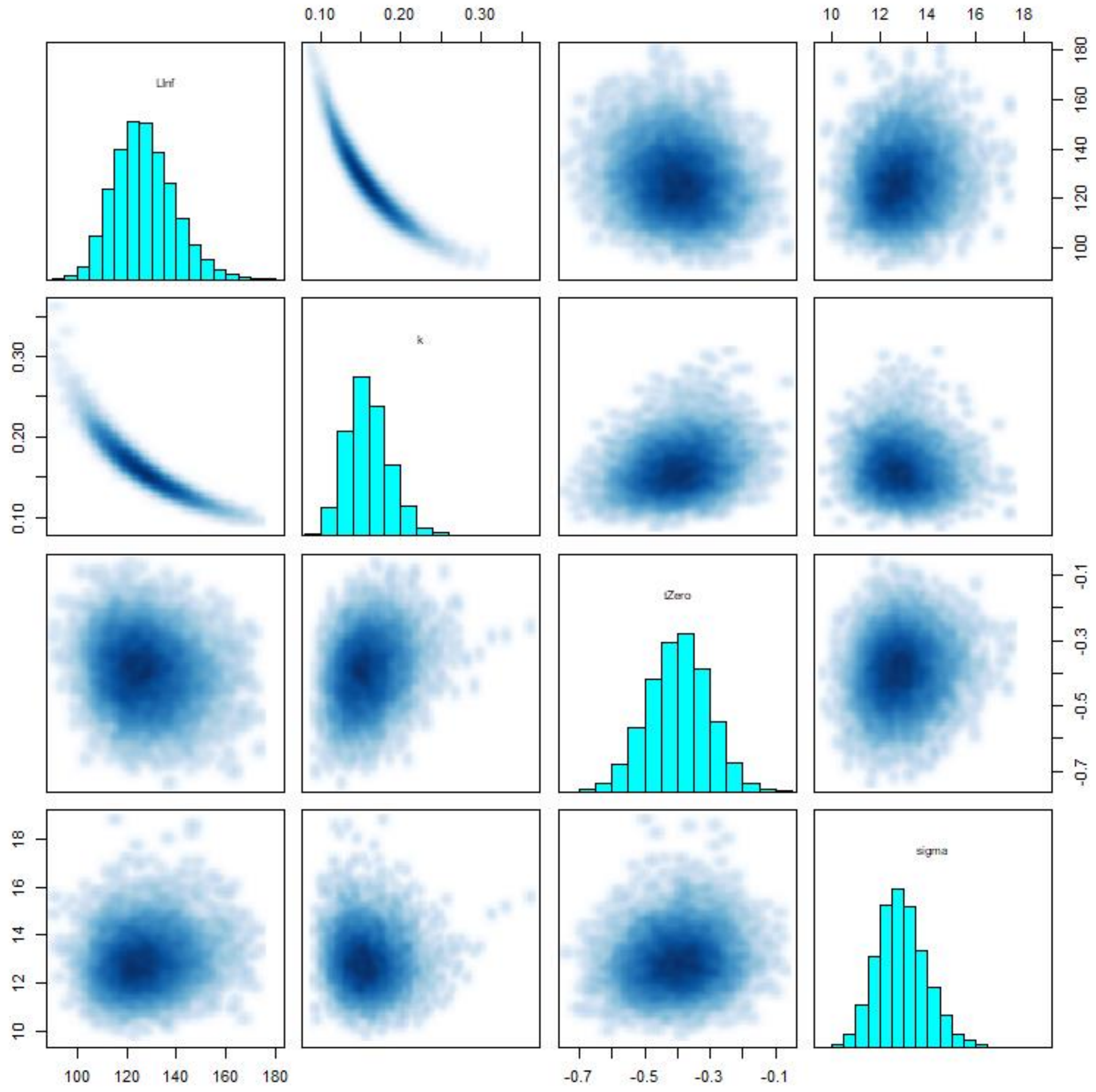


Figure S9: Pairs plot for the scale age model, showing the interactions between variables used in the model. The inherent negative correlation between $L_\infty$ and *k* is apparent by the presence of multiplicative non-identifiabilities (top left corner). Remaining parameters show even distributions in their respective bivariate plots as well as normally distributed univariate histograms, indicating no issues of colinearity between variables is present.

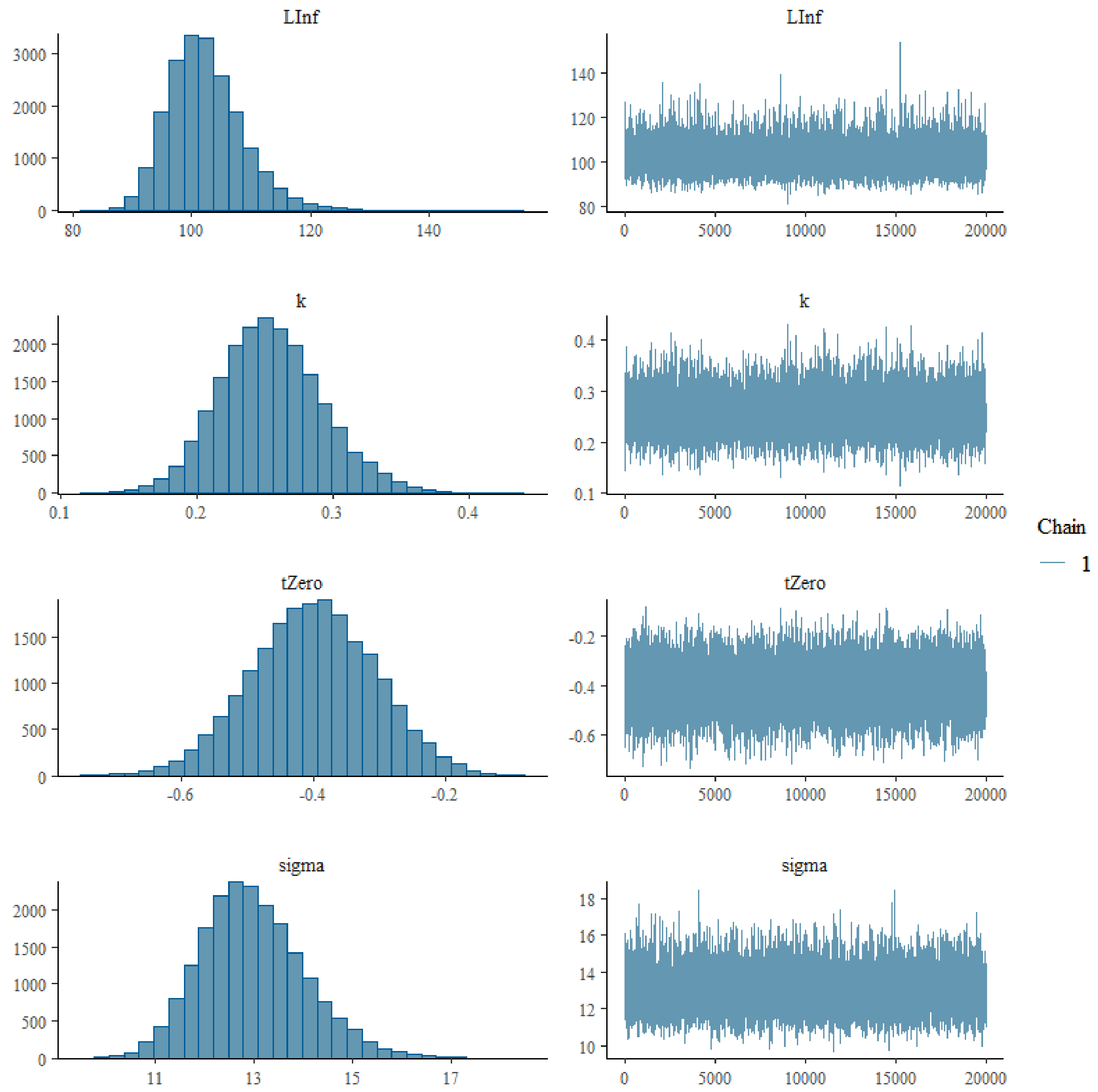


Figure S10: Trace and histogram plot for the corroborated age growth model. Traceplots (right side) indicate chains have mixed for all parameters and model efficiency was satisfactory. Parameter histograms (left side) indicate no multi-modality.

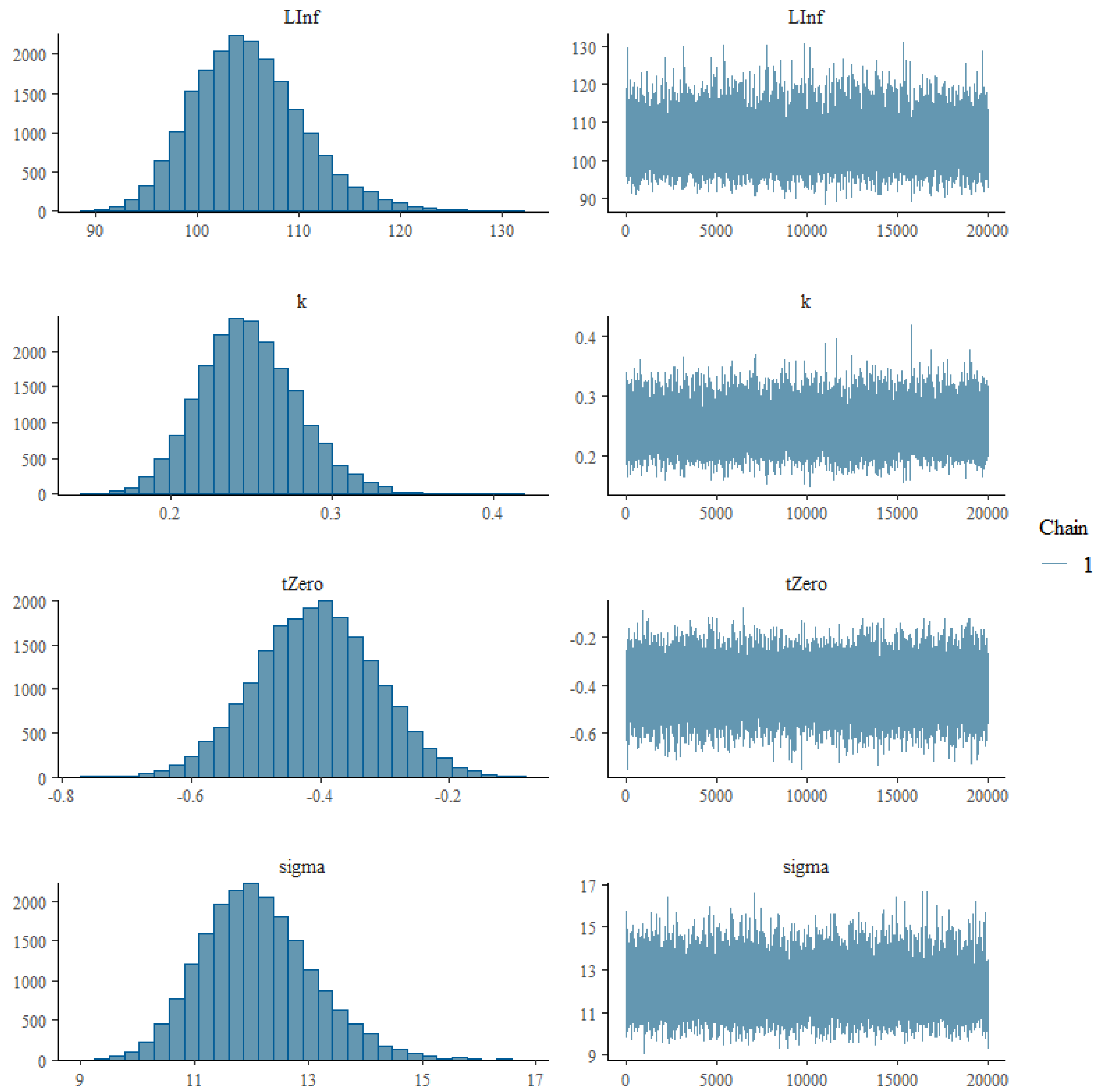


Figure S11: Trace and histogram plot for the otolith age growth model. Traceplots (right side) indicate chains have mixed for all parameters and model efficiency was satisfactory. Parameter histograms (left side) indicate no multi-modality.

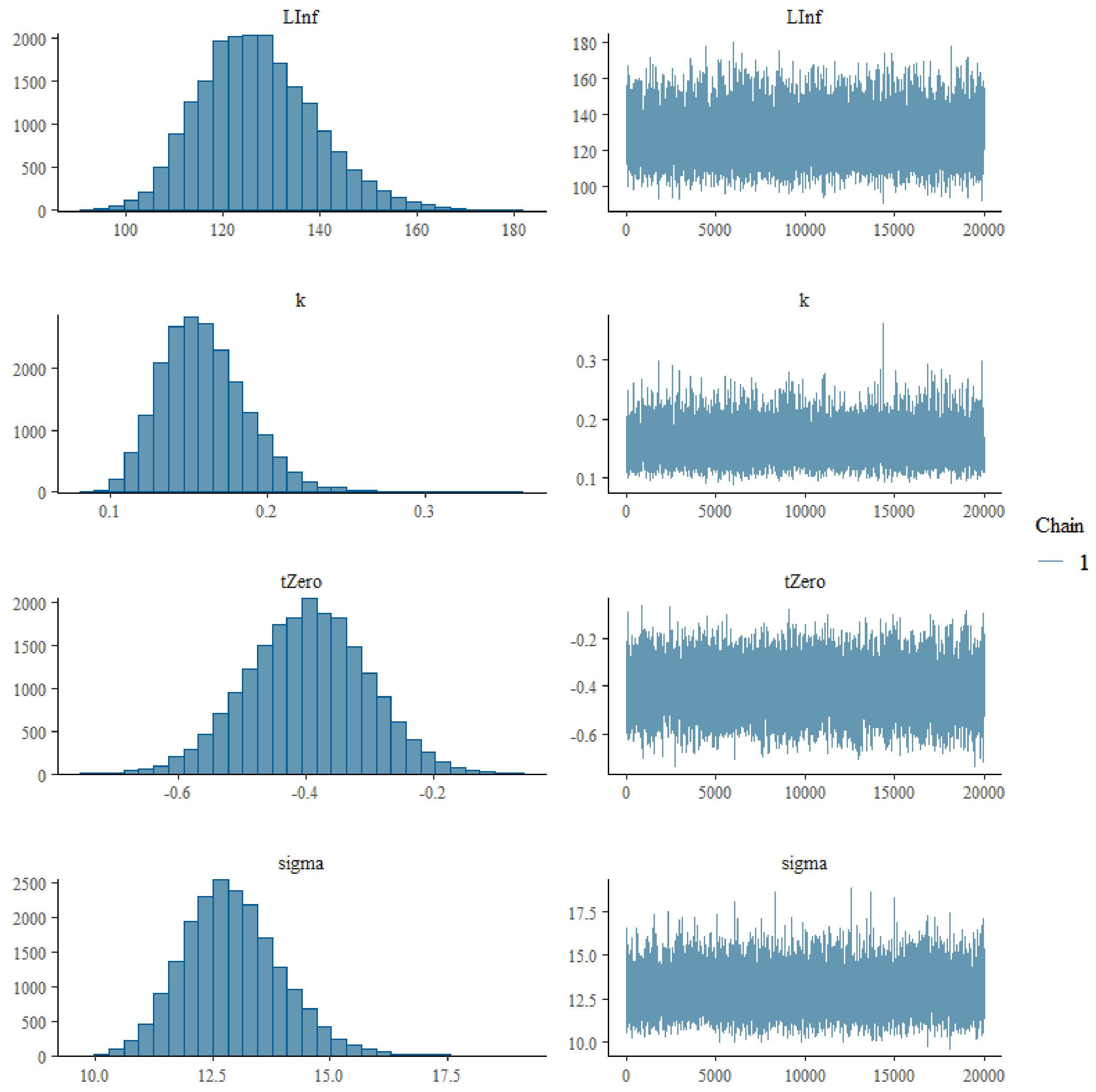

Figure S12: Trace and histogram plot for the otolith age growth model. Traceplots (right side) indicate chains have mixed for all parameters and model efficiency was satisfactory. Parameter histograms (left side) indicate no multi-modality.

**D: Parameter posterior credibility intervals for structure-specific growth models**

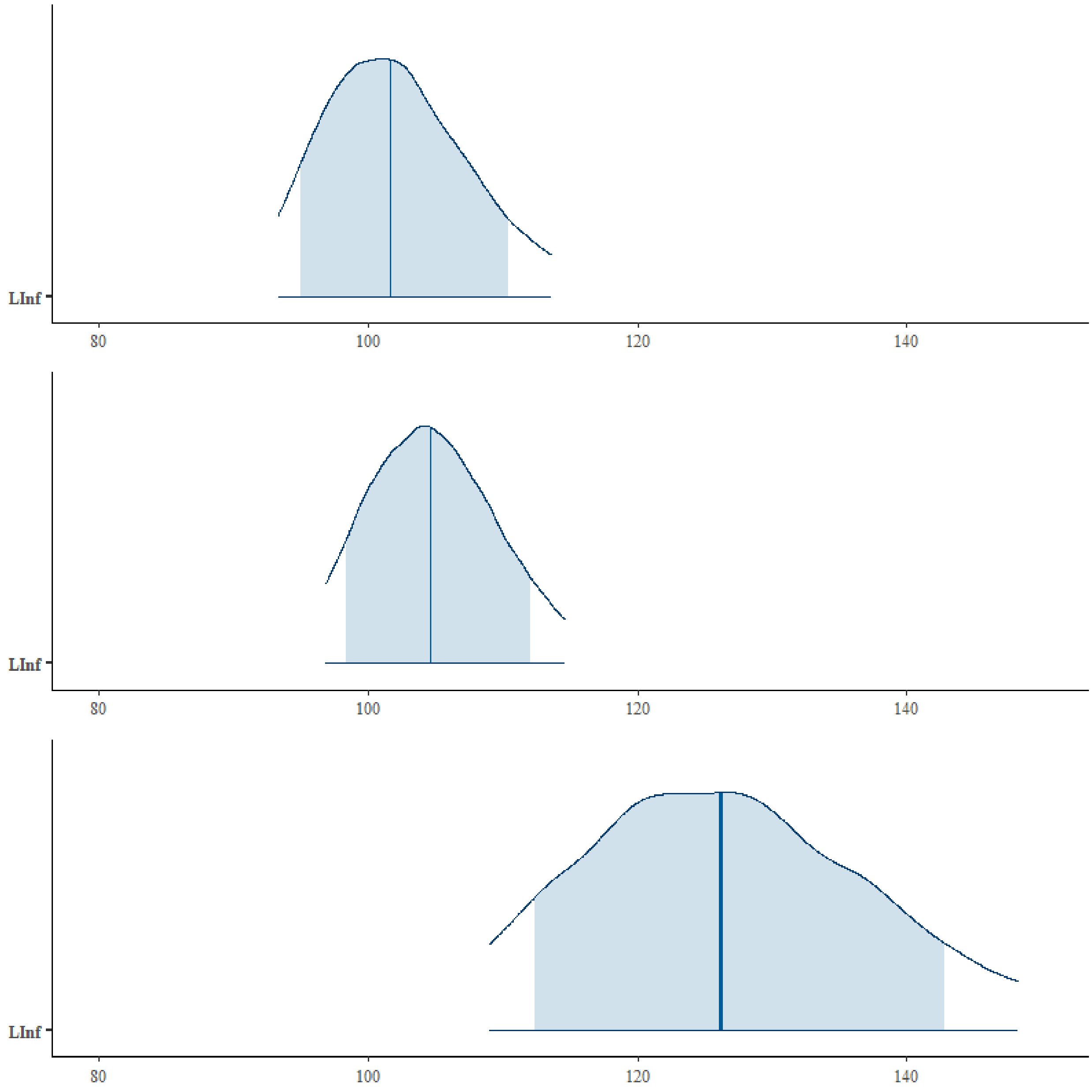


Figure S13: MCMC areas plot for the parameter $L_\infty$ of three structure growth models. The area outlined by the solid blue line represents the outer 90% credibility intervals of the posterior parameter distribution. The area shaded in light blue represents the inner 80% credibility interval of the posterior parameter distribution. The vertical blue lines represent the median point of the posterior parameter distribution.

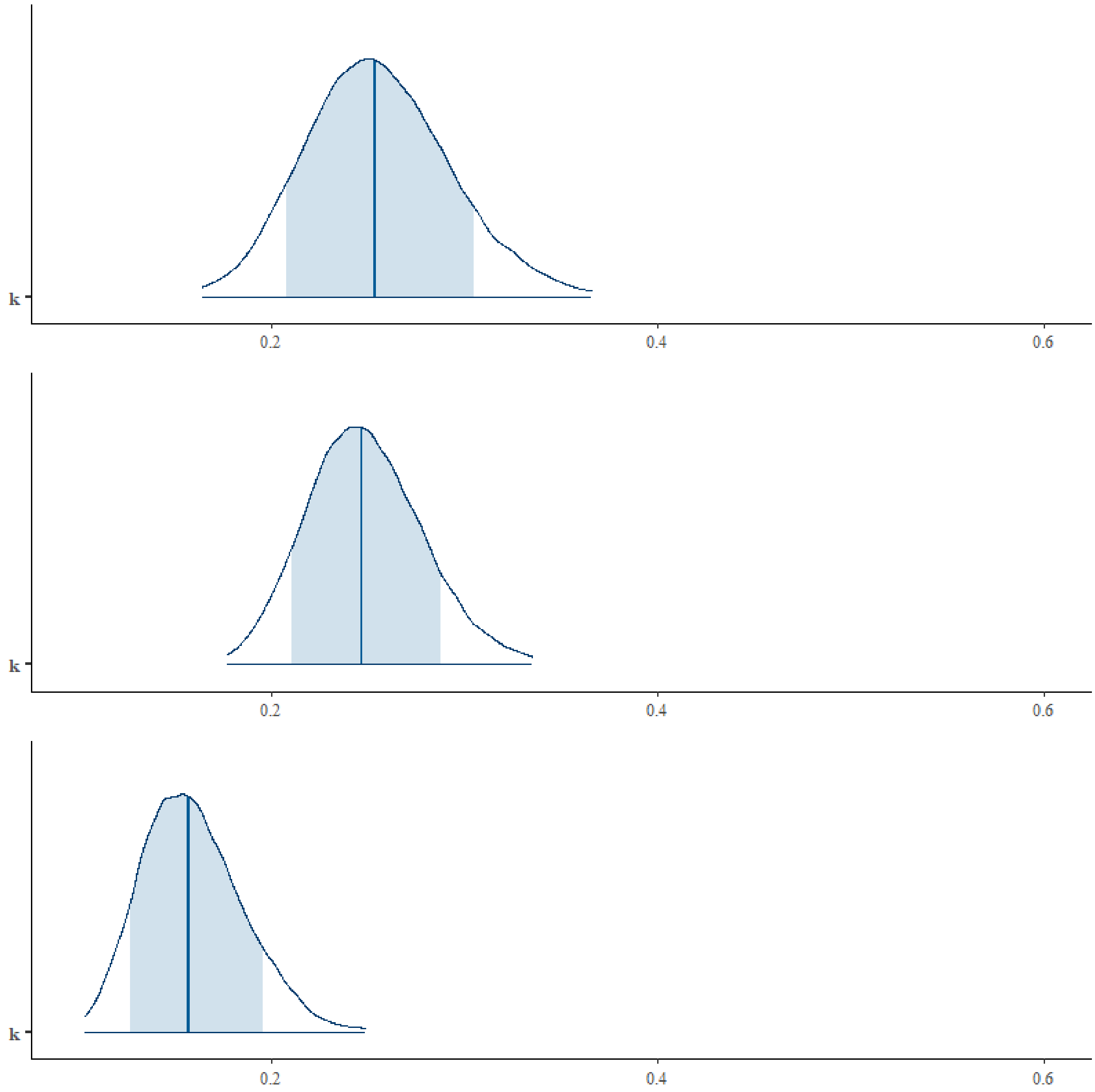


Figure S14: MCMC areas plot for the parameter *k* of three structure growth models. The area outlined by the solid blue line represents the outer 90% credibility intervals of the posterior parameter distribution. The area shaded in light blue represents the inner 80% credibility interval of the posterior parameter distribution. The vertical blue lines represent the median point of the posterior parameter distribution.

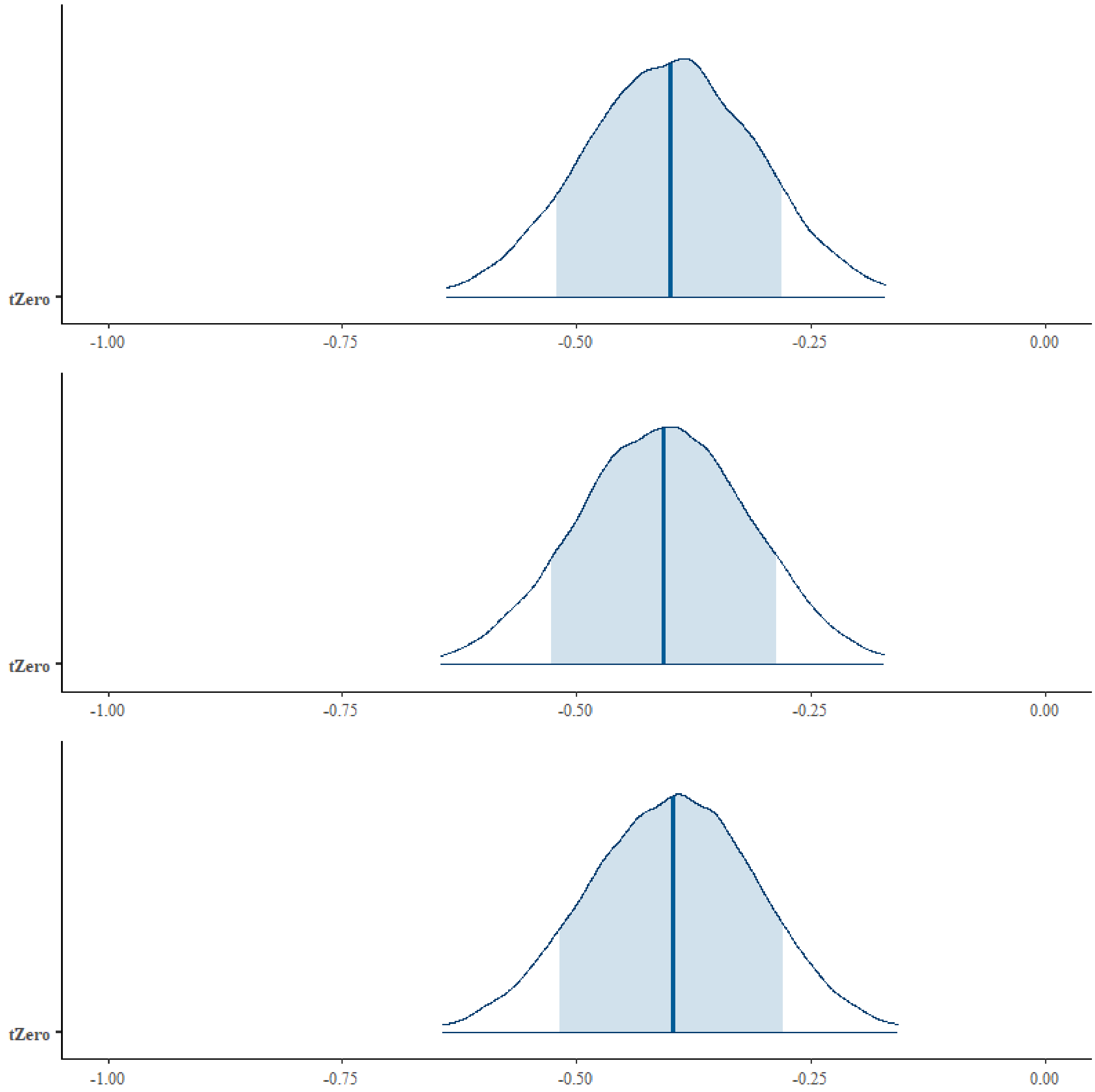


Figure S15: MCMC areas plot for the parameter $t_0$ of three structure growth models. The area outlined by the solid blue line represents the outer 90% credibility intervals of the posterior parameter distribution. The area shaded in light blue represents the inner 80% credibility interval of the posterior parameter distribution. The vertical blue lines represent the median point of the posterior parameter distribution.

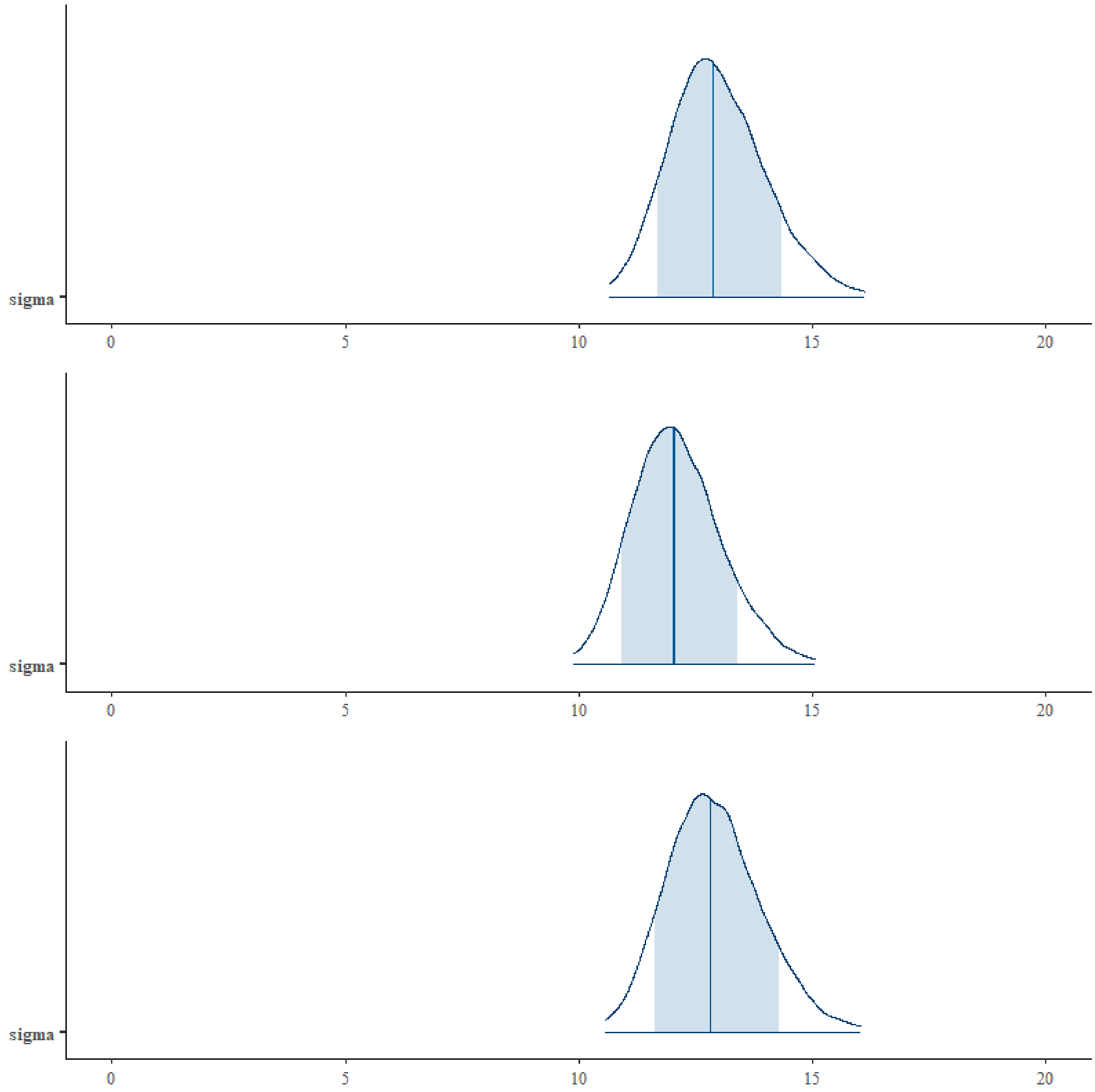


Figure S16: MCMC areas plot for the parameter σ of three structure growth models. The area outlined by the solid blue line represents the outer 90% credibility intervals of the posterior parameter distribution. The area shaded in light blue represents the inner 80% credibility interval of the posterior parameter distribution. The vertical blue lines represent the median point of the posterior parameter distribution.

**E: Expected $\delta^{18}O$ values in otolith aragonite**

Linear mixed effects models showed the predicted $\delta^{18}O$-values of both equation 1 and equation 2 to be significantly correlated to water temperature and $\delta^{18}O_{Water}$ (Table S1). Maximum likelihood ratio tests were used to test the significance of fixed effects. The models were fitted with 'lmer' using restricted maximum likelihood (REML) estimation in the lme4 package (Bates et al., 2015). A Gaussian error distribution was assumed, which was confirmed by visual inspection of the residuals and QQ plots. The random effect of month was found to have a significant effect, but it explained only marginal portions of the variance in the models (< 0.01%). Using the average deviation of $\delta^{18}O_{Water}$ (sd = 0.7 ‰) over the year at a constant temperature to model $\delta^{18}O_{Otolith}$ accounted for a variation of 0.7 ‰ in the predicted $\delta^{18}O_{Otolith}$ for both equations, whereas using the average deviation of water temperatures (sd = 6.5 °C) at constant $\delta^{18}O_{Water}$ to model $\delta^{18}O_{Otolith}$ accounted for a variation of 1.4 ‰ in $\delta^{18}O_{Otolith}$.

Table S1: Summary of fixed and random effects on the predicted $\delta^{18}O_{Otolith}$-values predicted by the two equations used for the calculation of theoretical $\delta^{18}O_{Otolith}$-values in a pike otolith.

| **Prediction equation 1 (Patterson et al., 1993)** | | | | **Prediction equation 2 (Geffen et al., 2012)** | | | |
|---|---|---|---|---|---|---|---|
| *Fixed effect* | *Estimates* | *CI* | *p* | *Fixed effect* | *Estimates* | *CI* | *p* |
| $\delta^{18}O_{Water}$ | 1 | 1.00 – 1.00 | **< 2.2e-16 *** ** | $\delta^{18}O_{Water}$ | 1 | 1.00 – 1.00 | **< 2.2e-16 *** ** |
| Temperature | -0.23 | -0.23 – -0.23 | **< 2.2e-16 *** ** | Temperature | -0.2 | -0.20 – -0.19 | **< 2.2e-16 *** ** |
| **Random Effects** | | | | **Random Effects** | | | |
| $\sigma^2$ | 2.76e-07 | | | $\sigma^2$ | 2.05e-07 | | |
| $\tau_{00\ month}$ | 8.76e-04 | | | $\tau_{00\ month}$ | 6.44e-04 | | |
| ICC | 1 | | | ICC | 0.99968219 | | |
| $N_{month}$ | 12 | | | $N_{month}$ | 12 | | |
| Observations | 24 | | | Observations | 24 | | |
| Marginal $R^2$ / Conditional $R^2$ | 0.999 / 1.000 | | | Marginal $R^2$ / Conditional $R^2$ | 0.999 / 1.000 | | |

**F: Assessing fisheries management reference points and optimal size limits**

Table S2: Equations of the age-structured pike population model and model output metrices.

| | Equation | Description |
|---|---|---|
| **3** | $L_a = L_\infty \left(1 - e^{-k(a-t_0)}\right)$ | Average length of an individual of age $a$ following the von Bertalanffy growth equation |
| **4** | $w_a = \alpha_w \, {L_a}^{\beta_w}$ | Length-weight relationship |
| **5** | $g(L_a, l_{low}, l_{up}) = \int_{l_{low}}^{l_{up}} \left(\sqrt{2\pi}\,\sigma\right)^{-1} \exp\left[-\frac{(l-L_a)^2}{2\sigma^2}\right] dl$ with $\sigma = L_a \cdot cv$ | Proportion of individuals of age $a$ with a length between $l_{low}$ and $l_{up}$, assuming a normally distributed intracohort variation in length around $L_a$ |
| **6** | $m_a = g(L_a, l_{mat}, \infty)$ | Proportion mature |
| **7** | $f_a = \alpha_f \, {w_a}^{\beta_f}$ | Weight-dependent fecundity |
| **8** | $R_{t+1} = \alpha_R \, E_t \, e^{-\beta_R \, N_{spawners,t}}$ | Age 1 recruits at time $t+1$ based on a Ricker-type stock recruitment function |
| **9** | $E_t = \sum_a N_{a,t} \, m_a \, f_a$ | Total number of eggs produced at time $t$ |
| **10** | $N_{spawners,t} = \sum_a N_{a,t} \, m_a$ | Spawning stock size at time $t$ which reduces egg survival by cannibalism |
| **11** | $\alpha_R = \frac{CR}{\sum_a \zeta_a \, m_a \, f_a}$ | Maximum egg survival (= effect of density-independent mortality on recruitment) |
| **12** | $\beta_R = \frac{\ln(CR)}{R_0 \sum_a \zeta_a \, m_a}$ | Density-dependent effect on egg survival (= effect of density-dependent mortality on recruitment) |
| **13** | $M_a = M \left(\frac{L_\infty}{L_a}\right)^{\vartheta}$ | Size-dependent natural mortality following the Lorenzen equation |
| **14** | $\zeta_{a+1} = e^{-M_a} \, \zeta_a \qquad \vert \zeta_1 = 1$ | Survivorship unfished |
| **15** | $v_a^c = g(L_a, l_{\min c}, l_{\max c})$ | Selectivity of capture |
| **16** | $v_a^r = g(L_a, l_{\min r}, l_{\max r})$ | Selectivity of retention |
| **17** | $p_a = \frac{v_a^r}{v_a^c}$ | Proportion of capture retained |
| **18** | $H_a = F \, v_a^c \, p_a$ | Harvest mortality |
| **19** | $D_a = F \, v_a^c \, (1 - p_a) \, d$ | Discard mortality |
| **20** | $Z_a = M_a + H_a + D_a$ | Total mortality |
| **21** | $N_{a+1,t+1} = N_{a,t} \, e^{-Z_a}$ | Numbers over time |
| **22** | $Y_{B,t} = \sum_a \frac{H_a}{Z_a} N_{a,t} \, (1 - e^{-Z_a}) \, w_a$ | Yield (biomass harvested) at time $t$ |
| **23** | $B_t = \sum_a N_{a,t} \, w_a$ | Biomass at time t |
| **24** | $Y_{N,t} = \sum_a \frac{H_a}{Z_a} N_{a,t} \, (1 - e^{-Z_a})$ | Number of harvested individuals at time $t$ |
| **25** | $N_{vuln,t} = \sum_a N_{a,t} \, v_a^c$ | Number of individuals vulnerable to catch in population at time $t$ |
| **26** | $N_{trophy,t} = \sum_a N_{a,t} \, p_a^{trophy}$ with $p_a^{trophy} = g(L_a, l_{trophy}, \infty)$ | Number of trophy fish in population at time t |
| **27** | $U_t = \ln\left(\frac{Y_{B,t}}{MSY_B} \cdot 100\right) + \ln\left(\frac{Y_{N,t}}{MSY_N} \cdot 100\right) + \ln\left(\frac{B_t}{B_0} \cdot 100\right) + \ln\left(\frac{N_{vuln,t}}{N_{vuln,0}} \cdot 100\right) + \ln\left(\frac{N_{trophy,t}}{N_{trophy,0}} \cdot 100\right)$ | Log utility function including different output metrices, each scaled relative to its maximum value (t=0 refers to the unfished state, $MSY_B$ represents the classic maximum sustainable biomass yield and $MSY_N$ is similar, but for yield in terms of number harvested). |

Table S3: Symbols, meanings, values, units and sources of model parameters. Frost and Kipling (1967), Myers et al. (1999).

| Symbol | Meaning | Value | Unit | Data source |
|---|---|---|---|---|
| $L_\infty$ | Average asymptotic length | Varied within 90 % CI estimated for each aging method | $cm$ | Own data |
| $k$ | Van Bertalanffy growth coefficient | Varied within 90 % CI estimated for each aging method | $year^{-1}$ | Own data |
| $t_0$ | Theoretical age at which length of 0 | Varied within 90 % CI estimated for each aging method | $year$ | Own data |
| $cv$ | Coefficient of variation in VBGF | 0.13 | | Frost & Kipling 1967 |
| $\alpha_w$ | Scaling parameter for length-weight relationship | 0.0045 | For converting $cm$ into $g$ | Own data |
| $\beta_w$ | Allometric parameter of length-weight relationship | 3.107 | For converting $cm$ into $g$ | Own data |
| $M$ | Minimum adult instantaneous natural mortality (at large sizes) | 0.25 | $year^{-1}$ | Assumed |
| $\vartheta$ | Lorenzen size-dependent mortality power | 0.5 | | Ahrens et al. 2020 |
| $a_{max}$ | Maximum age | 15 | $year$ | Own data |
| $l_{min\ c}$ | Minimum length at which fully vulnerable to capture | 40.0 | $cm$ | Own data |
| $l_{max\ c}$ | Maximum length at which fully vulnerable to capture | 150.0 | $cm$ | Set above $L_\infty$ |
| $l_{min\ r}$ | Minimum length of retention (= Minimum-length limit) | 50.0 (varied in scenarios) | $cm$ | Current harvest regulation |
| $l_{max\ r}$ | Maximum length of retention (i.e., upper bound of harvest slot or maximum length limit) | 150.0 (varied in scenarios) | $cm$ | Set above $L_\infty$ as currently no harvest slot |
| $F$ | Total instantaneous fishing mortality on individuals with a selectivity of 1 | 0.2 (varied in scenarios) | $year^{-1}$ | van Gemert et al. 2021 |
| $d$ | Discard mortality (proportion of individuals dying after release) | 0.078 | | Hühn & Arlinghaus 2011 |
| $l_{mat}$ | Length at maturation (females) | 37.5 | $cm$ | Own data |
| $\alpha_f$ | Scaling parameter of weight - fecundity relationship | 9.8 | | Own data |
| $\beta_f$ | Power parameter relating fecundity to weight | 1.12 | | Own data |
| $R_0$ | Unfished equilibrium total number of age-1 recruits | $1.0 \cdot 10^6$ | | Set at a value that generates biomasses and yields within the observed range |
| $CR$ | Goodyear compensation ratio | 6.1 | | Myers et al. 1999 |

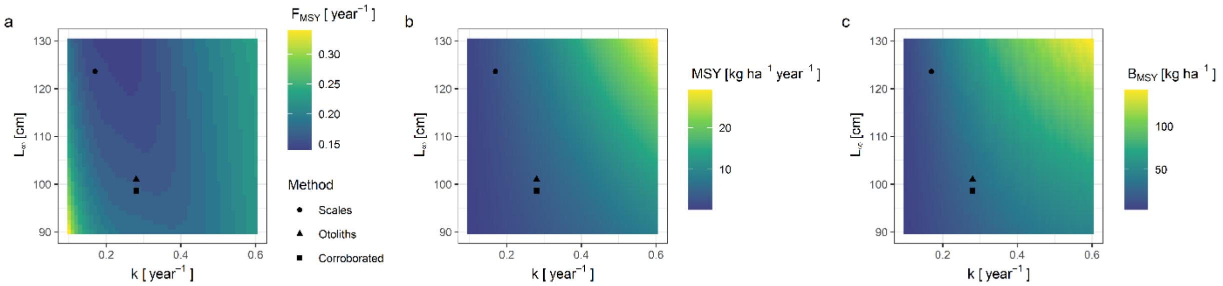


Figure S17: Model predictions of fishery reference points a) $F_{MSY}$, b) $MSY$ and c) $B_{MSY}$ depending on the growth coefficient $k$ and asymptotic length $L_\infty$ systematically changed over a broad range of values, assuming a fixed theoretical age at length 0 ($t_0 = -0.4\ years$). The black markers indicate the mean values of $k$ and $L_\infty$ obtained from the different aging methods. It is important to note that $k$ and $L_\infty$ were changed independently from other life history traits. This is biologically not correct given well-known life-history relationships between $k$ and natural mortality, for example, but was done to illustrate the pure effect of bias in growth parameter estimation.

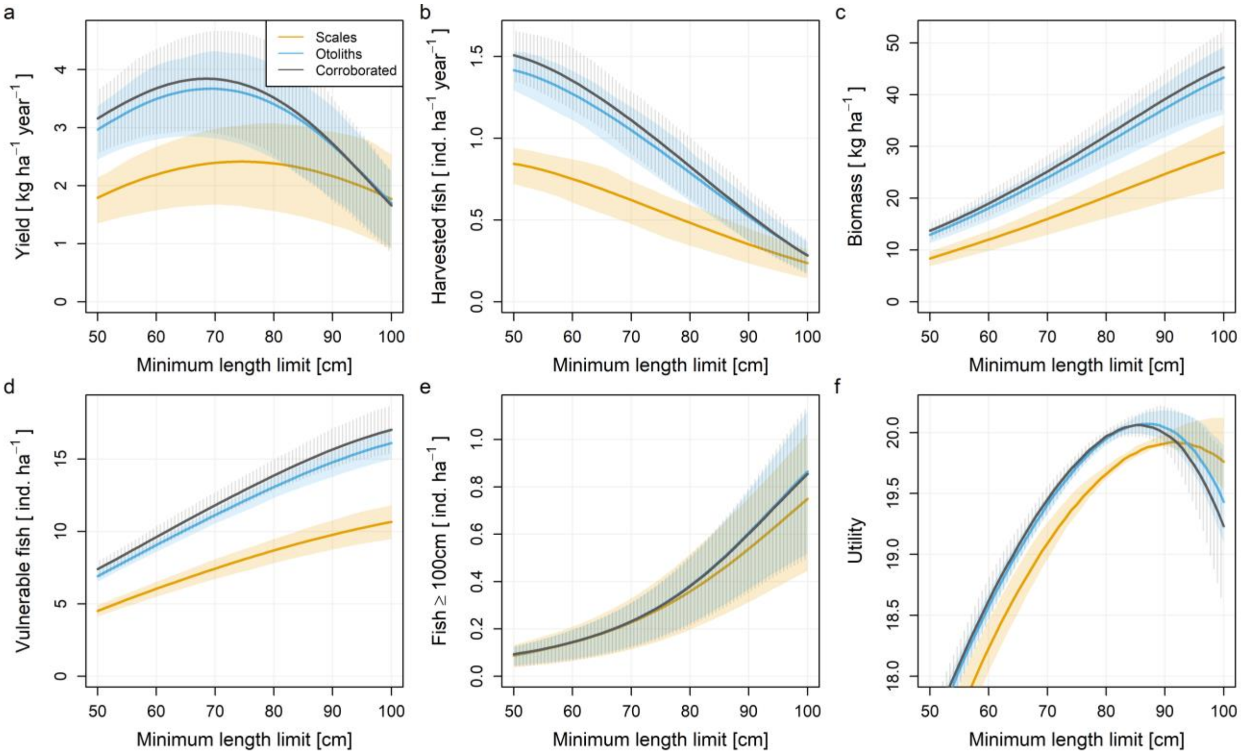


Figure S18: Effect of minimum length limit (currently 50 cm) on the output of the model, parametrized with different growth curves based on scales, otoliths and corroborated age, for an increased instantaneous fishing mortality of 0.4 $years^{-1}$ (standard scenario shown in main text: 0.2 $years^{-1}$). The output metrices include a) harvested biomass, b) number of harvested fish, c) stock biomass, d) density of fish vulnerable to catch, e) density of trophy pike and f) utility which includes the five aforenamed metrices. The lines represent mean values of 100 simulation replicates with different growth parameters for each aging method, randomly sampled from normal distribution within the 90%-credibility interval estimated from the growth data. The shaded area equals the respective interquartile range.

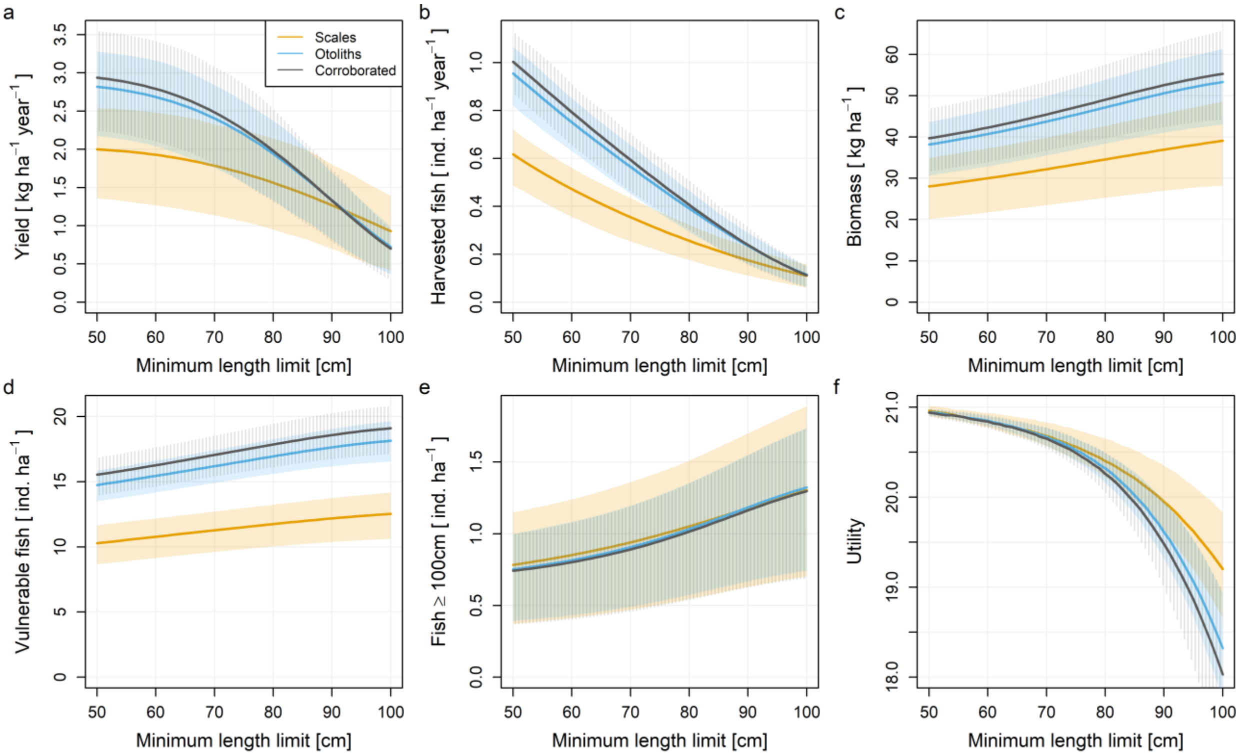


Figure S19: Effect of minimum length limit (currently 50 cm) on the output of the model, parametrized with different growth curves based on scales, otoliths and corroborated age, for a decreased instantaneous fishing mortality of 0.1 $years^{-1}$ (standard scenario shown in main text: 0.2 $years^{-1}$). The output metrices include a) harvested biomass, b) number of harvested fish, c) stock biomass, d) density of fish vulnerable to catch, e) density of trophy pike and f) utility which includes the five aforenamed metrices. The lines represent mean values of 100 simulation replicates with different growth parameters for each aging method, randomly sampled from normal distribution within the 90%-credibility interval estimated from the growth data. The shaded area equals the respective interquartile range.

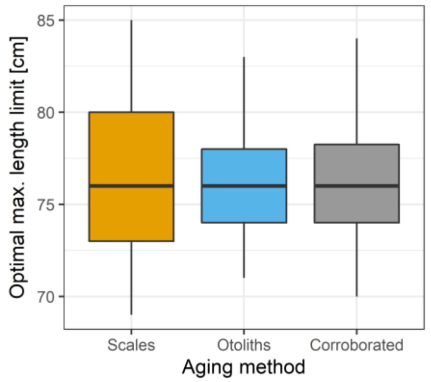


Figure S20: Boxplot showing the optimal maximum length limit (i.e. the upper limit of the harvest slow with the lower limit fixed at 50 cm) predicted by the model for different growth curve parametrizations based on three different ageing methods. Variation in optimal maximum length limit results from randomly sampling growth parameters from normal distribution within the 90%-credibility interval obtained from growth data for each aging method. Outliers were not plotted.

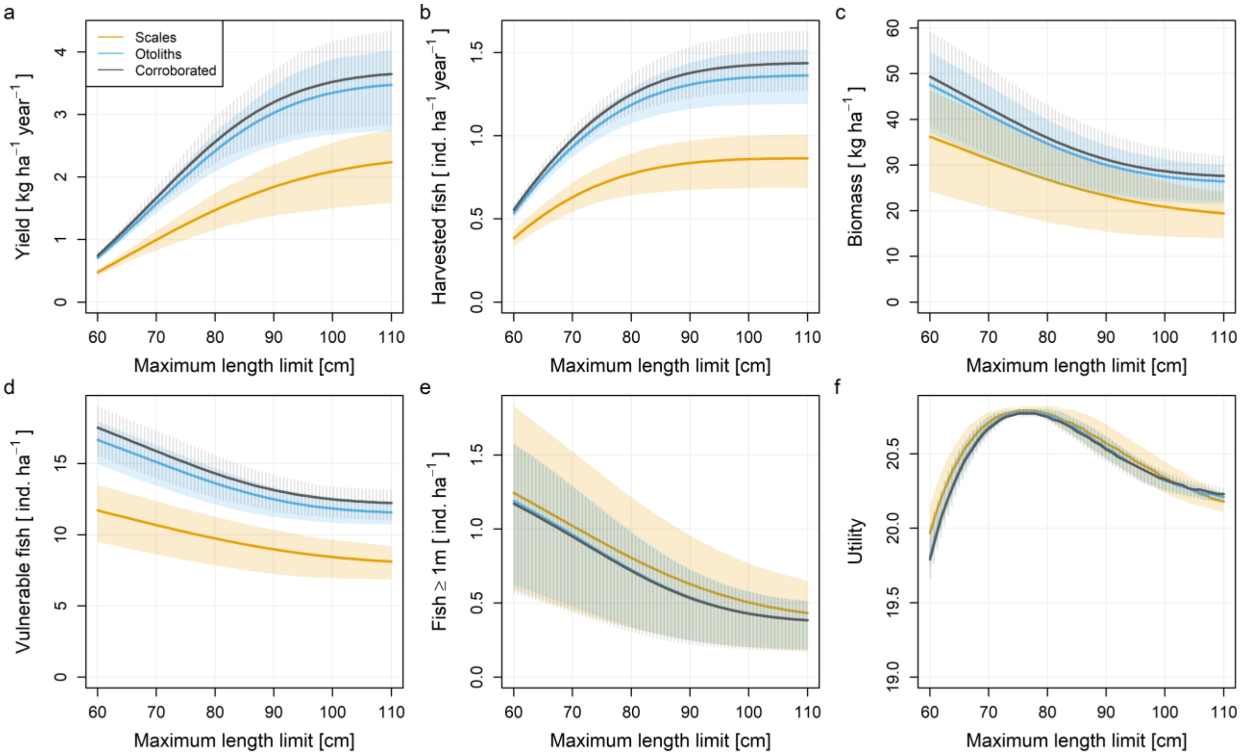


Figure S21: Effect of maximum length limit (i.e. upper limit of harvest slot with lower limit fixed at 50 cm) on the output of the model, parametrized with different growth curves based on scales, otoliths and corroborated age. The output metrices include a) harvested biomass, b) number of harvested fish, c) stock biomass, d) density of fish vulnerable to catch, e) density of trophy pike and f) utility which includes the five aforenamed metrices. The lines represent mean values of 100 simulation replicates with different growth parameters for each aging method, randomly sampled from normal distribution within the 90%-credibility interval estimated from the growth data. The shaded area equals the respective interquartile range.
